# A detailed Molecular Network Map and Model of the NLRP3 Inflammasome

**DOI:** 10.1101/2023.05.31.543045

**Authors:** Marcus Krantz, Daniel Eklund, Eva Särndahl, Alexander Hedbrant, the X-HiDE consortium

## Abstract

The NLRP3 inflammasome is a key regulator of inflammation that responds to a broad range of stimuli. The exact mechanism of activation has not been determined, but there is a consensus on cellular potassium efflux as a major common denominator. Once NLRP3 is activated, it forms high-order complexes together with NEK7 that trigger aggregation of ASC into specks. Typically, there is only one speck per cell, consistent with the proposal that specks form – or end up at – the centrosome. ASC polymerisation in turn triggers caspase-1 activation, leading to maturation and release of IL-1β and pyroptosis, i.e., highly inflammatory cell death. Several gain-of-function mutations in the NLRP3 inflammasome have been suggested to induce spontaneous activation of NLRP3 and hence contribute to development and disease severity in numerous autoinflammatory and autoimmune diseases. Consequently, the NLRP3 inflammasome is of significant clinical interest, and recent attention has drastically improved our insight in the range of involved triggers and mechanisms of signal transduction. However, despite recent progress in knowledge, a clear and comprehensive overview of how these mechanisms interplay to shape the system level function is missing from the literature. Here, we provide such an overview as a resource to researchers working in or entering the field, as well as a computational model that allows for evaluating and explaining the function of the NLRP3 inflammasome system from the current molecular knowledge. We present a detailed reconstruction of the molecular network surrounding the NLRP3 inflammasome, which account for each specific reaction and the known regulatory constraints on each event as well as the mechanisms of drug action and impact of genetics when known. Furthermore, an executable model from this network reconstruction is generated with the aim to be used to explain NLRP3 activation from priming and activation to the maturation and release of IL-1β and IL-18. Finally, we test this detailed mechanistic model against data on the effect of different modes of inhibition of NLRP3 assembly. While the exact mechanisms of NLRP3 activation remains elusive, the literature indicates that the different stimuli converge on a single activation mechanism that is additionally controlled by distinct (positive or negative) priming and licensing events through covalent modifications of the NLRP3 molecule. Taken together, we present a compilation of the literature knowledge on the molecular mechanisms on NLRP3 activation, a detailed mechanistic model of NLRP3 activation, and explore the convergence of diverse NLRP3 activation stimuli into a single input mechanism.

## Introduction

The innate immune system serves as an immediate and essential defence towards exogenous and endogenous threats. Its activity is influenced by a number of pattern-recognition receptors (PRRs) that recognize pathogen-and damage-associated molecular patterns (PAMPs/DAMPs). The PAMPs are families of highly conserved molecular patterns that are indicative of (evolutionarily) common pathogens, while the DAMPs are endogenous compounds that are released from cells or the extracellular matrix during tissue or cell damage, or exogenous molecules, such as air pollution particles. PAMP and DAMPs can be recognised at the plasma membrane, in the endolysosomal system, or in the cytoplasm. Inflammasomes belong to the latter group, by being centred around intracellular receptors that can nucleate the formation of a large intracellular protein complex [1]. When activated, these receptors trigger inflammasome assembly and thereby caspase-1 activation, proteolytic processing and release of cytokines, such as interleukin (IL)-1β and IL-18, and eventually pyroptosis – a highly pro-inflammatory form of cell death – via gasdermin D-dependent pore formation. Following Gasdermin D pore formation and cell death, plasma membrane rupture is mediated by NINJ1; another pore forming protein that is assembled by a so far undetermined mechanism [2, 3]. Inflammasome activation has also been linked to direct anti-microbial actions through autophagy, presumably to clear the infection [4], and the direct bactericidal effect of mature and active gasdermin D [5]. Hence, inflammasomes are critical immune regulators at the intersection between immediate antimicrobial defence and intercellular signalling.

The NLRP3 inflammasome is the odd one out of the inflammasome sensors, as it responds to a large repertoire of signals. Instead of recognising a conserved molecular pattern, NLRP3 responds to a wide range of triggers that at a first glance have little in common. These include perturbation to the membrane potential with ionophoric toxins, such as nigericin, perturbation to mitochondrial function, and exposure to crystals and crystal forming compounds. In response to each of these triggers, NLRP3 oligomers serve as a nucleation centre for ASC polymerisation. ASC is recruited to NLRP3 through homotypic PYD domain interactions and can continue to polymerise, forming a large ASC speck that can be monitored through e.g. fluorescence microscopy. These ASC specks in turn recruit pro-caspase-1, resulting in its activation through proximity-induced trans-autoproteolysis. Active caspase-1 cleaves pro-IL-1β and pro-IL-18 into their mature and active forms, as well as gasdermin D, which allows the latter to form pores in the plasma membrane to release IL-1β and IL-18 [6], and, eventually, to trigger pyroptosis. However, none of these triggers suffice to activate the NLRP3 inflammasome in a healthy system, as NLRP3 needs to be licensed for activation by a priming signal (reviewed in e.g.: [7], [8]). Experimentally, LPS is usually used as a priming signal, which, through TLR4, Tak1, IKKβ, and NFκB affects NLRP3 at two distinct levels: through induction of gene transcription and by posttranslational licensing. Hence, activation of NLRP3 requires two or three steps, depending on whether the cell type shows a high basal expression of NLRP3 or not, including: 1) (conditionally) transcriptional priming, 2) posttranslational licensing, and 3) activation by a trigger. This three-step picture is complicated further by the apparently wide range and partially unrelated trigger effects, as well as by the fact that multiple signalling pathways control licensing through multiple modification sites in different components of the NLRP3 inflammasome. Consequently, both the nature of activation and the integration of licensing signals remain opaque.

In this work, we take a systems biology approach to gain understanding on the NLRP3 inflammasome mechanisms. We perform an in-depth literature review and curation to compile the existing mechanistic knowledge on NLRP3 regulation into a formal knowledge base. Briefly, we use an established workflow for reconstruction of signal transduction networks, which relies on iterative literature curation, network validation and gap-filling [9]. The goal is to provide a comprehensive mechanistic model, i.e., a model that includes all relevant components and processes of the system under study, and which describes those as an unbroken link of mechanisms and causalities from system input to output. To this end, we use rxncon, the reaction-contingency language, to formalise the network in terms of elemental reactions and contingencies (see methods for details)[10, 11]. Elemental reactions represent minimal and decontextualised reaction events, similar to the reaction centre in rule-based modelling [12]. Contingencies represent the regulatory constraints on reactions, similar to the reaction context in rule-based modelling. The resulting (rxncon) knowledge base has been processed by the rxncon toolbox to visualise the network and to allow parameter-free simulation [13]. In particular, we have made use of the scalable regulatory graph to visualise the information flow through the network (Figures 2-12)[10], to make the content of the knowledge base easily accessible, and to help the reconstruction process. Building such models is an excellent way to add value to existing data. The fundamental idea is to collect the available knowledge of NLRP3 activation in a single knowledge resource, much like the biochemical pathways maps of metabolism, and to analyse how far the current molecular level knowledge of NLRP3 activation can explain what is known about the system level function. In this particular case, the question raised is to what extent the molecular knowledge of NLRP3 inflammasome signalling can explain the release of mature cytokines in response to the inputs that are known to trigger this release, and to what extent the effect of known mutations or drug perturbations on this response can be reproduced in the model. Clearly, this work has a strong component of literature review. However, it also has a number of features that go further. First, the statements in a mechanistic model need to be concrete and precise, and these explicit interpretations of the data can be evaluated individually. Second, the model must be internally consistent, which means that all apparent contradictions must be resolved. Third, the model can be tested through simulation, to make sure that the expected system level function (activation or not) emerges from the assembled molecular mechanisms. Fourth, the annotated model with individual references for each model entry increase reusability, making it easy for the community to update and extend the model as new knowledge becomes available. Hence, the model constitutes an explicit and internally consistent compilation of the current molecular level knowledge that is complemented with graphs that visualise these molecular processes in detail. We envisage it as a research community resource to serve as an entry point for novices in the field, as a guide for future experiments, and as a contribution to the discussion on what really activates NLRP3.

## Material and Methods

### Literature curation and network reconstruction

The network reconstruction was performed with a previously developed and demonstrated workflow [9, 14, 15]. Literature curation started from a number of reviews, essentially providing an initial “parts list” in terms of components, processes and signals that can prime or trigger NLRP3. This starting point helped guide a targeted literature search in two complementary directions: First, on specific interactions, modifications, and mechanisms that constitute the actual signal transduction, to map out the actual signal transduction processes. Second, on the different trigger signals and their connection to NLRP3 activation, to determine if and how they could be attributed to a common cellular perturbation and how that perturbation could be sensed by NLRP3. PubMed, Google and perplexity.ai were used complementarily to search for literature relevant for specific questions, and these papers, together with references in initial reviews and retrieved papers, were used to build the network model. The network reconstruction was compiled in the second generation rxncon language [11] as elemental reactions and contingencies (see below). Each elemental reaction and contingency is associated with one or more references through PubMed IDs, or clearly marked as a model hypothesis (see column “!Reference:Identifiers:pubmed” in Suppelementary Table 1). The following papers are referenced in the model: [8, 16-70].

**Table 1:** NLRP3 inflammasome triggers by trigger class. The triggers can be divided into three general categories: (I) ionophoric compounds and compounds triggering ion fluxes indirectly (such as extracellular ATP), (II) compounds destabilising the lysosomes, and (III) inhibitors of the mitochondrial electron transport chain (ETC). Most, but not all, triggers can be suppressed by high extracellular KCl (column KCl, Y=Yes, can be suppressed, N=No, cannot be suppressed). Most lysosomal destabilising compounds require phagocytosis or endocytosis (P/E), with the exception of LLOMe. N/A = not applicable, blanks = no information found.

| **Table 1: NLRP3 inflammasome triggers by trigger class** | **KCl** | **P/E** | **Reference(s)** |
| --- | --- | --- | --- |
| **Type I (Ionophorous)** |  |  |  |
| Aerolysin, hemolysin and MARTX (*Aeromonas*) | Y | N/A | [83] |
| ATP | Y | N/A | [39, 84] |
| Biglycan |  | N/A | [85] |
| Exolysin (*Pseudomonas*) | Y | N/A | [86] |
| Hemolysin and non-hemolytic enterotoxin (*Bacillus*) | Y | N/A | [87, 88] |
| Hemolysin, M1 protein and pneumolysin (*Streptococcus*) | Y | N/A | [89-91] |
| Hemolysins (*Escherichia*) | Y | N/A | [92, 93] |
| Hemolysins and leukocidins (*Staphylococcus*) | Y | N/A | [94-98] |
| Hemolysins and MARTX (*Vibrio*) | Y | N/A | [99, 100] |
| Listeriolysin O (*Listeria*) | Y | N/A | [101] |
| Nigericin and valinomycin (*Streptomyces*) | Y | N/A | [18, 102, 103] |
| Perfringolysin O and tetanolysin O (*Clostridium*) | Y | N/A | [104, 105] |
| Serum amyloid A (SAA) |  | N/A | [106] |
| Shiga toxins (*Shigella*) | Y | N/A | [107] |
| **Type II (lysosomal destabilisation)** |  |  |  |
| Alum | Y | Y | [108, 109] |
| Amyloid-β (fibrillar) | | Y | [110] |
| Asbestos | Y | Y | [111] |
| Calcium pyrophosphate dihydrate crystals (CPPD) | Y | Y | [109, 112] |
| Carbon nanotubes | Y | Y | [113, 114] |
| Chitosan | Y | Y | [102, 115] |
| Cholesterol crystals | Y | Y | [116, 117] |
| Copper oxide nanoparticles |  | Y | [118] |
| Dextran sodium sulfate (DSS) | Y | Y | [119] |
| Graphene oxide |  | Y | [120] |
| Hyaluronan |  | Y | [121] |
| Leu-Leu-O-methyl ester (LLOMe) | Y | N/A | [64, 109] |
| Monosodium urate crystals (MSU) | Y | Y | [39, 108] |
| Oxidised LDL | Y | Y | [122] |
| Quartz/silica crystals | Y | Y | [108, 109] |
| **Type III (ETC inhibitors)** |  |  |  |
| CL097 | N | N/A | [39] |
| Imiquimod | N | N/A | [39] |

### The rxncon language and encoding of knowledge

The network reconstruction was compiled in the second generation rxncon language [11]. The rxncon network definition is based on site-specific elemental states. Importantly, elemental states capture the state at a single residue or domain, and multiple elemental states are typically needed to fully define the protein (micro)state (discussed in [71]). Elemental states do not correspond to disjunct entities, as components with multiple sites (binding domains or modification residues) are represented by multiple nodes in the graphs (e.g., each NLRP3 molecule may be phosphorylated on Ser5, Ser198, Ser295, Ser806, and/or Tyr861). The possible combinations of these states are not represented unless necessary in a contingency. E.g., AKT mediated phosphorylation of NLRP3 Ser5 requires AKT to be phosphorylated on both Thr308 and Ser473 (or bound to the activator SC79), and to be bound to phosphoinositide (PI; which in turn need to be 4-phosphorylated to bind AKT (Figure 1)). This makes the network representation more abstract than in microstate-based formalisms such as the process description diagram formalism [72], but brings three distinct advantages: First, the elemental state representation has an excellent congruence with empirical data, making network reconstruction precise and straightforward [71]. Second, the omission of unnecessary (in the perspective of empirical data) enumeration of state combinations abrogates the combinatorial complexity [73]. Third, the (regulatory graph) representation emphasises causality, providing a clear overview of the information flow through even very complex networks [10]. The rxncon network is defined at two complementary levels that are both defined in terms of elemental states and hence both correspond directly to empirical data: (1) Elemental reactions defines decontextualised reaction events. (2) Contingencies defines contextual constraints on elemental reactions, in terms of (Boolean combinations of) elemental states or inputs.

**Figure 1:**
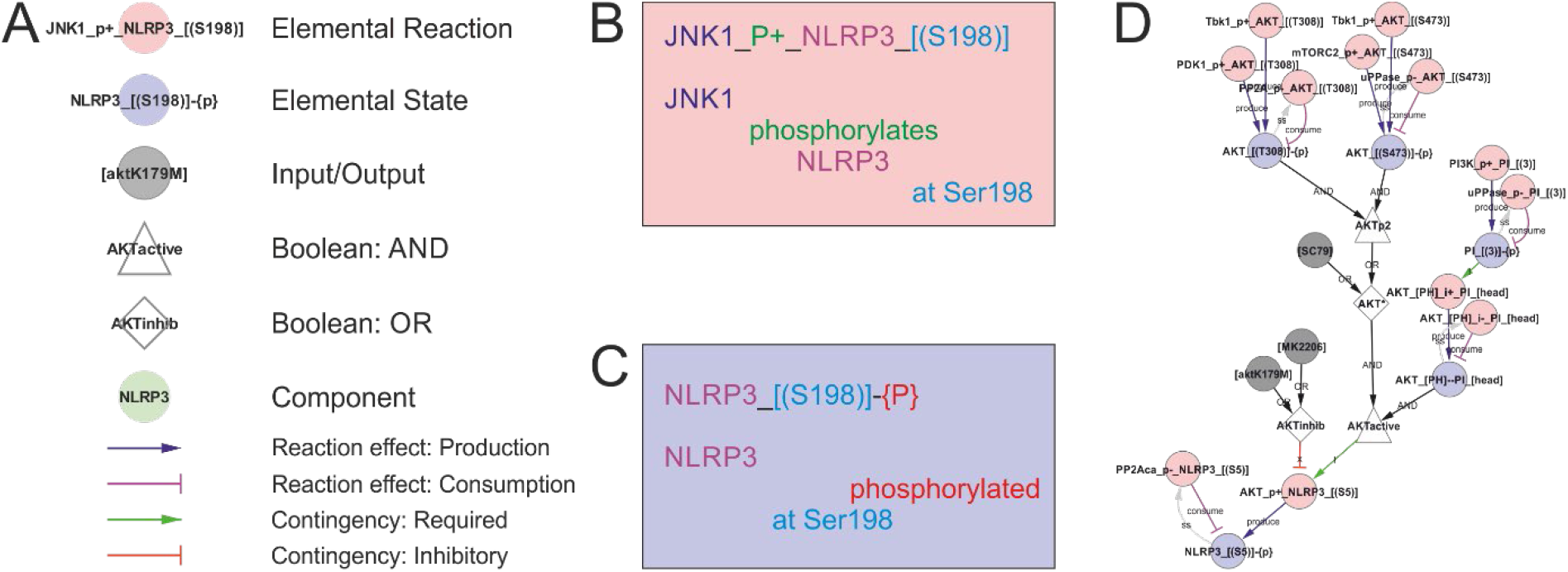
The rxncon regulatory graph format. The regulatory graph focusses on the causality in signal transduction. The information is encoded in elemental states (blue nodes e.g., phosphorylated state of protein x at site y), which are changed by, but also controls, elemental reactions (red nodes e.g., protein z phosphorylates protein x at site y). Reaction-to-state edges display the effect of elemental reactions (production/consumption of elemental states) and states-to-reaction edges display the regulatory effect of states on reactions. The information flow can be followed along the directed edges in the network, but the details of the reactions and states are encoded in the node labels. **(A) The principal node and edge types** in the NLRP3 network. **Nodes:** Red: Elemental reactions; Blue: Elemental states (Note that neutral elemental states (unmodified, unbound) are excluded from the visualisation); Grey: Inputs and Outputs (e.g., nigericin as an input, and IL-1β release as an output in NLRP3 activation); White: Boolean operators (Triangles: AND, Diamonds: OR, Octagons: NOT); Green: Components (e.g., a protein or mRNA). **Reaction-to-state edges:** Blue: Production; Magenta: Consumption; Black: Synthesis or Degradation. **State-to-reaction edges:** Green: State is required for the target reaction; Red: State inhibits the target reaction. **(B) The elemental reaction label** contains three elements: ComponentA (JNK1; in this case the catalyst), a reaction type (in this case a phosphorylation; P+), and ComponentB (NLRP3). Depending on reaction type, components may be defined with loci (in this case only for NLRP3; the residue Ser198). Loci are identified by hard brackets “[]” for interaction domains, and nested hard and normal brackets “[()]” for residues). **(C) The elemental state label** contains the Component(s) with locus/loci depending on state type. Here, the state corresponds to NLRP3 phosphorylated at Ser198. **(D) Example of a complex requirement**: The control of Ser5 phosphorylation state in NLRP3 (“NLRP3_[(S5)]-{P}”). This residue is phosphorylated (“P+”) by AKT (“AKT_P+_NLRP3_[(S5)]") and dephosphorylated (“P-“) by PP2Aca (“PP2Aca_P-_NLRP3_[(S5)]”). AKT phosphorylation requires that AKT is active (“AKTactive”) and it is blocked by AKT inhibition (“AKTinhib”). AKT activation and inhibition are defined by Boolean combination of states: AKTinhib is an OR gate of an AKT inhibitor (“MK2206”) and an AKT mutation (“AKTK179M”), meaning that either alone suffices to inhibit the reaction. “AKTactive” is an AND gate of PI-binding (“AKT_[PH]--PI_[head]”) and “AKT*”, meaning that both criteria must be fulfilled. Hence, both activation criteria must be TRUE, and both inhibition criteria FALSE, for the reaction to be active. “AKT*” is in turn an OR gate of chemical activation (“SC95”) or double phosphorylation of AKT at Thr308 and Ser473. Hence, (nested) Boolean combination of elemental states and/or inputs can specify arbitrarily detailed regulatory constraints. Finally, the phosphorylation status of AKT Thr308 and Ser473 are controlled by phosphorylation by Tbk1 (e.g. “Tbk1_P+_AKT_[(T308)]"), PDK1 (only Thr308), and mTORC2 (only Ser473) and by dephosphorylation (“P-“).

Elemental state are site specific states in one or two components. For example, the Ser5 phosphorylation is NLRP3_[(S5)]-{P}, consisting of a component name (NLRP3), a locus (residue serine 5: _[(S5)]) and a modification (Phosphate: -{P}). Correspondingly, the bond with NEK7 is NEK7_[clobe]- -NLRP3_[HD2LRR], where the first protein (NEK7) via the clobe-domain (_[clobe]) binds (--) the second component (NLRP3) via the HD2-LRR region of NLRP3 (_[HD2LRR]). Importantly, an elemental state gives no information on the state at any other residue or domain.

Elemental reactions include one or two components, but are always defined through two components (A and B; which in monomolecular reactions are the same). Each component is defined at a certain resolution (Component, Domain, or Residue), depending on the reaction type and the component’s role in the reaction, as described in detail elsewhere [11]. The elemental reactions contain no information on the states of the components beyond the state(s) that change through the reaction. For example, the phosphorylation of Ser5 by AKT (AKT_P+_NLRP3_[(S5)]) requires the site to be unmodified (NLRP3_[(S5)]-{0}; where “-{0}” indicates an “empty” residue), but have no other intrinsic requirements. The contextual states, i.e., those that do not change through the reaction (i.e., the requirement for AKT phosphorylation and PI binding for activation), are defined as contingencies. The elemental reactions are defined in the ReactionList sheet of Supplementary Table 1.

The contingencies define the regulatory constraints on reactions. The qualitative model presented here includes three types of contingencies: Required (!), inhibitory (x), and no effect (0). Simple constraints may be defined by a single contingency, e.g. that caspase-1 processing (Caspase1_cut_Caspase1_[(pro)]) require (!) caspase-1 dimerisation (Caspase1_[card1]-- Caspase1_[card2]). However, more complex requirements can also be accommodated by (possibly nested) Boolean combinations (AND, OR, or NOT) of Elemental states and inputs, as illustrated by AKT activation above. These Booleans are also defined in the contingency list, where the <BOOLEAN> (identified by “<>”) appears both as a target – on the lines where the <BOOLEAN> is defined – and as modifier for elemental reactions and outputs, and when it is part of the definition of another Boolean expression (in nested Booleans). Finally, the contingency list is used to define system [Inputs] and [Outputs] (identified by “[]”), which constitutes the boundary of the system. The contingencies are defined in the ContingencyList sheet of Supplementary Table 1

In this model, we make a special use of inputs and outputs to describe Signal 2. The effects that trigger NLRP3 cannot be efficiently described at the level of single molecules, hence we use placeholder entities that are connected as a chain of inputs/outputs that control each other. While it would have sufficed to include the most downstream inputs, centrosomal PI(4)P or CL, for the model to be functional, the inclusion of the steps – even in this relatively crude format – allows us to elaborate on the hypothesis and distinguish different type of inputs – e.g. in term of potassium efflux and suppression by extracellular KCl. In addition, we deem the graph useful as an overview complementing the discussion in the text.

For a detailed description of the rxncon language, see:. [10, 11]

### Visualisation, model generation and simulation

The generation of the rxncon regulatory graph for visualisation and the bipartite Boolean model (bBM) for simulation was performed with the rxncon2regulatorygraph.py script and rxncon2boolnet script, respectively. Both scripts are part of the rxncon toolbox that can be downloaded from GitHub (https://github.com/rxncon/rxncon; without dependencies), installed from the python package index through “pip install rxncon”, or through kboolnet [74]. Simulation was performed using BoolNet as described previously [13, 15]: Starting from a highly artificial initial state, the model was simulated until it reached its natural off state. From this state, it was exposed to different treatment by setting the corresponding inputs (grey circles in the regulatory graph) to True. The signal transmission through the network and the effect on the outputs were monitored to determine if the NLRP3 inflammasome was activated or not.

## Results

### The reconstruction process and scope

The reconstruction process was based on manual literature curation. Starting from an overview, based on several review articles, targeted literature searches were used to clarify the relationships between components and the importance of specific modifications and bonds, as well as to connect the apparently unrelated triggers to a common mechanism of activation. The starting point is taken from the basic assumption that the molecular functions are the same across cell types, and that any difference between (isogenic) cells can be explained by expression differences rather than differences in molecular function. Inclusion of data from different organisms (primarily mouse in addition to human cells and cell lines) is less straightforward but has been used as indicated in the model. Importantly, as a mechanistic model requires direct mechanistic connections between components, components and functions can only be included when their mechanistic function in the network is known. Concretely, this means that for a molecule to be added into the model it is not enough that it has been shown to interact with e.g. NLRP3; there must also be a known functional outcome of that interaction. For example, the model does not include cathepsin B, thioredoxin-interacting protein (TXNIP), or caspase-8, despite their reported roles in NLRP3 regulation. TXNIP, which dissociates from its partner thioredoxin upon oxidative stress and elevated ROS, has been suggested as a binding partner to NLRP3 after dissociation [75]. In silico modelling of predicted binding has indicated conformational changes in the pyrin domain of NLRP3 by TXNIP binding, facilitating interactions with ASC [76]. However, TXNIP has been found to be dispensable for NLRP3 activation by ATP, MSU, and islet amyloid polypeptide [8], and subsequent knock-out studies have been inconclusive. Thus, TXNIP has not been included in the model. Also, cathepsin B has been excluded due to inconclusive literature, and studies that have shown it to be dispensable for NLRP3 activation (reviewed in [77]). Similarly, the model does not include caspase-8, as we have not found any mechanistic information on how it is connected to NLRP3 activation. Hence, as all caspase-8 dependent mechanisms are absent from the model, the model is effectively casp8-/-, and consequently NEK7 is absolutely required for NLRP3 activation [78]. However, it is important to bear in mind that some of the activation seen in experimental data may be due to the caspase-8 dependent, NEK7 independent, mechanism that is not included in the model. Other components have additional functions beyond those covered in the model. E.g., Tbk1/IKKe are involved in controlling NLRP3 phosphorylation at Ser5, but also have an impact even when this site is mutated [55]. However, as the mechanism of the latter effect remains unknown, this effect is not accounted for in the model. The complete knowledge base can be found in Supplementary Table 1, and it is visualised as a rxncon regulatory graph in Figure 2 and Supplementary Figure 1. For clarity, the network is divided and discussed in terms of modules, which are presented in figures 3-12. Below, the knowledge, interpretation, and implementation in the model will be described in detail.

**Figure 2:**
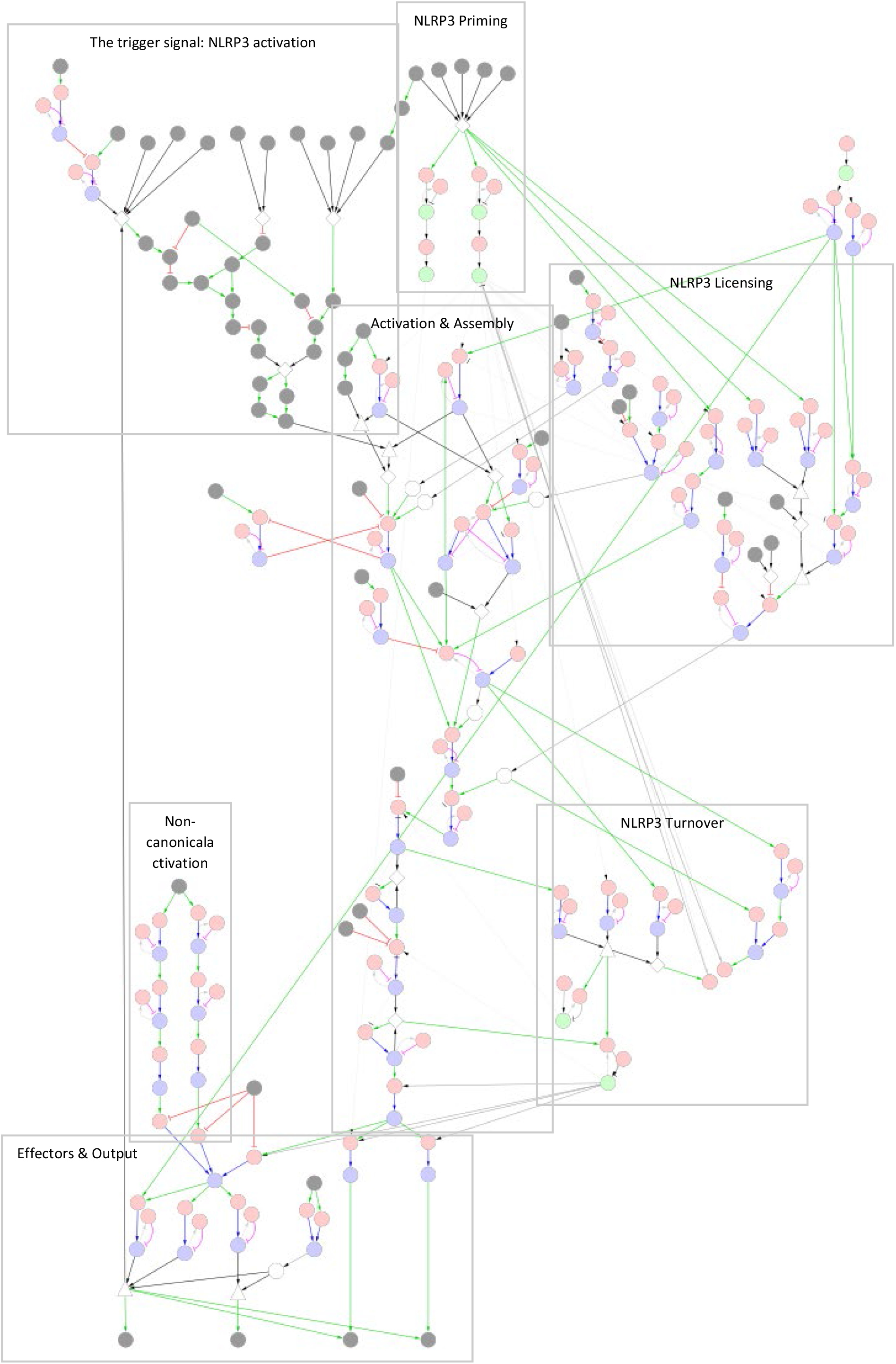
The NLRP3 inflammasome network. The NLRP3 inflammasome is activated by two signals: Signal 1, which primes and licenses the system for activation, and Signal 2, which triggers assembly and activation of the NLRP3 inflammasome. Legend: Nodes represent elemental reactions (red), elemental states (blue), components (green), and inputs/outputs (grey). Reaction-to-state edges represent the effect of the reaction on the target state (production/synthesis create states/components, consumption/degradation remove states/components). State-to-reaction edges represent the regulatory effect of the state on the target reaction (! = required for reaction, x = blocks reaction). Grey edges indicate that components or states participate in reactions. More complex requirements are defined as AND or OR combinations of states (and/or inputs), and are represented as white triangles and diamonds, respectively. Negation of states (NOT) are represented by white octagons. Note that neutral (unmodified, unbound) states are excluded from the regulatory graph for clarity. A high-resolution version with labels is included as Supplementary Figure 1.

**Figure 3:**
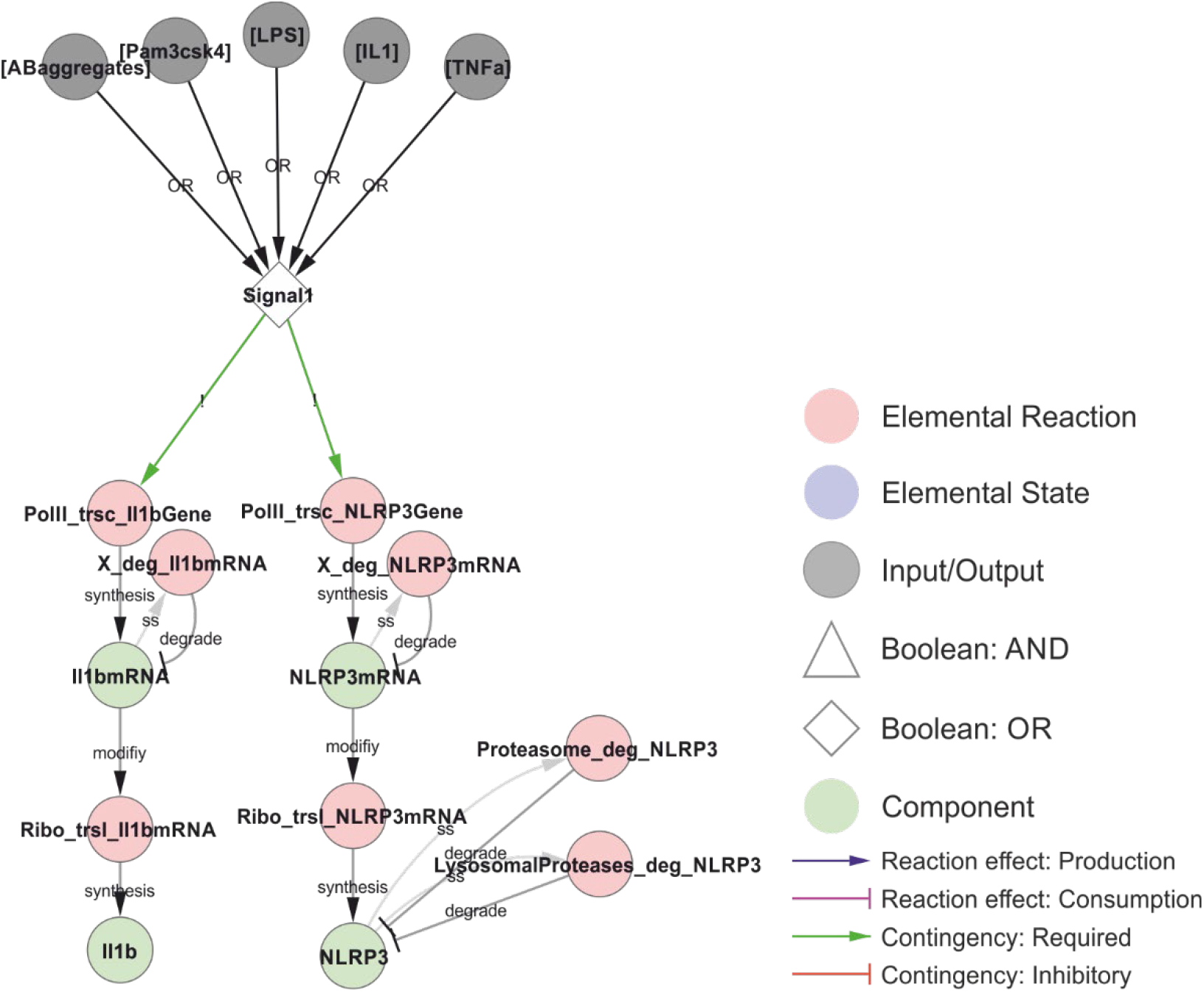
Transcriptional priming of NLRP3 and pro-IL-1β. The common denominator between the priming and licensing signals (Signal 1) is the activation of NFκB (omitted in the model), resulting in the transcription and translation of NLRP3 and pro-IL-1β. The mRNA is turned over (placeholder reactions catalysed by “X”), and NLRP3 is targeted for either proteasomal or lysosomal degradation depending on ubiquitylation (see below). The priming signals considered in this model are: LPS (Lipopolysaccharides; signalling via TLR4), Pam3csk4 (a synthetic triacylated lipopeptide; signalling via TLR1/2), IL-1 (interleukin-1, signalling via IL-1R1), TNF (tumour necrosis factor, signalling via TNFR1/2) and AB-aggregates (amyloid-β aggregates; through toll-like receptors). *Walkthrough: Either of five stimuli (grey nodes) can constitute Signal 1 (OR-gate), which is required for transcription (red nodes) of IL-1β (left) and NLRP3 (right). Transcription produces IL-1β mRNA and NLRP3 mRNA (green nodes) which in turn are translated (red nodes) to produce IL-1β and NLRP3. Transcripts and the NLRP3 protein are also degraded (red nodes). The model does not consider IL-1β degradation*.

### NLRP3 priming through NFκB-mediated transcription

The NLRP3 inflammasome is activated in three steps: transcriptional priming, posttranslational licensing, and triggering. Priming and licensing (also referred to as posttranslational priming) are induced by “Signal 1”, which corresponds to exposure to PAMPs (e.g. LPS (TLR4 ligand), Pam3CSK4 (TLR1/2 ligand)), or cytokines (e.g. IL-1 or TNF), which activate signalling through Tak1, IKKα/β, and NFκB [28, 79]. The model also includes amyloid-β aggregates, which have been found to act both as Signal 1 and, after endocytosis, as Signal 2 [27]. Signal 1 leads to the transcriptional induction of NLRP3 and pro-IL-1β (Figure 3). The priming event is necessary for inflammasome activation in some cell types, while others, such as human monocytes [80], have sufficiently high basal expression of NLRP3 and only need (posttranslational) licensing to enable a trigger to activate the NLRP3 inflammasome. In the model, this is implemented as a direct dependence of NLRP3 and pro-IL-1β transcription on Signal 1.

### NLRP3 licensing through posttranslational modification

NLRP3 inflammasome activation is also controlled by posttranslational licensing. Effectively, it constrains NLRP3 activation in space and time, to ensure activation only at the right place at the right time. NLRP3 activation is restricted by several posttranslational modifications that are only partially dependent on the priming signal. The picture is somewhat complicated by the observation that overexpression of NLRP3 is sometimes sufficient to override the need of licensing, as for example in HeLa cells [34]. This may indicate that licensing only increases the probability of activation, but that activation by sheer numbers is possible even without a licensing signal. It may also indicate that NLRP3 is licensed through release from a negative modification or stoichiometric inhibition, i.e. that one or more modifications or interaction partners keeps NLRP3 in an inactive state, and that the (limited) capacity of this inactivation is exhausted upon overexpression.

NLRP3 contains many potential modification sites. The model includes phosphorylation of Ser5, Ser198, Ser295, Ser806, and Tyr861, as the functional impact of these are mechanistically well characterised. These phosphorylation sites are spread across all three main domains of NLRP3: the PYD domain (residues 1-134), NACHT domain (residues 135-649; including the FISNA (residues 135-217) and the nucleotide binding domain (NBD)), and the LRR domain (residues 650-1036) [44]. In addition, the model accounts for K48-linked ubiquitylation of K496 – which targets NLRP3 for proteasomal degradation – and K63-linked ubiquitylation of the LRR domain – which prevents self-association and hence activation of NLRP3, and which targets NLRP3 for autophagy and hence lysosomal degradation. Apart from the control of self-association, ubiquitylation will be discussed further under NLRP3 turnover below.

The most well understood licensing modification of NLRP3 is indeed negative (Figure 4). Phosphorylation of Ser5 in the pyrin domain (PYD) of NLRP3 inhibits homotypic PYD-PYD interactions and therefore prevents PYD polymerisation, which is required for ASC recruitment and polymerisation [58]. Ser5 is phosphorylated by AKT [42] and dephosphorylated by PP2Acα [58]. AKT is considered constitutive [42] and activated by phosphorylation on Thr308 (by PDK1) and Ser473 (by mTORC2) [65], and by binding to PI(3,4)P2 or PI(3,4,5)P3 [20]. Hence, this constitutive activity is likely to be spatially restricted to membrane containing these phosphoinositides, i.e., primarily the plasma membrane and early endosomes [20]. This suggests that Ser5 phosphorylation imposes a spatial restriction on NLRP3 activation, consistent with the lack of ASC speck formation at the plasma membrane. However, AKT has also been shown to be activated by Tbk1/IKKε in a (LPS-) priming dependent fashion [48], in which the priming signal leads to a transient Tbk1/IKKε activation that delays and/or reduces the assembly of the NLRP3 inflammasome and hence IL-1β release [55]. Phosphorylation of Ser5 also stabilises NLRP3 [42], suggesting that activation leads to increased turnover of NLRP3.

**Figure 4:**
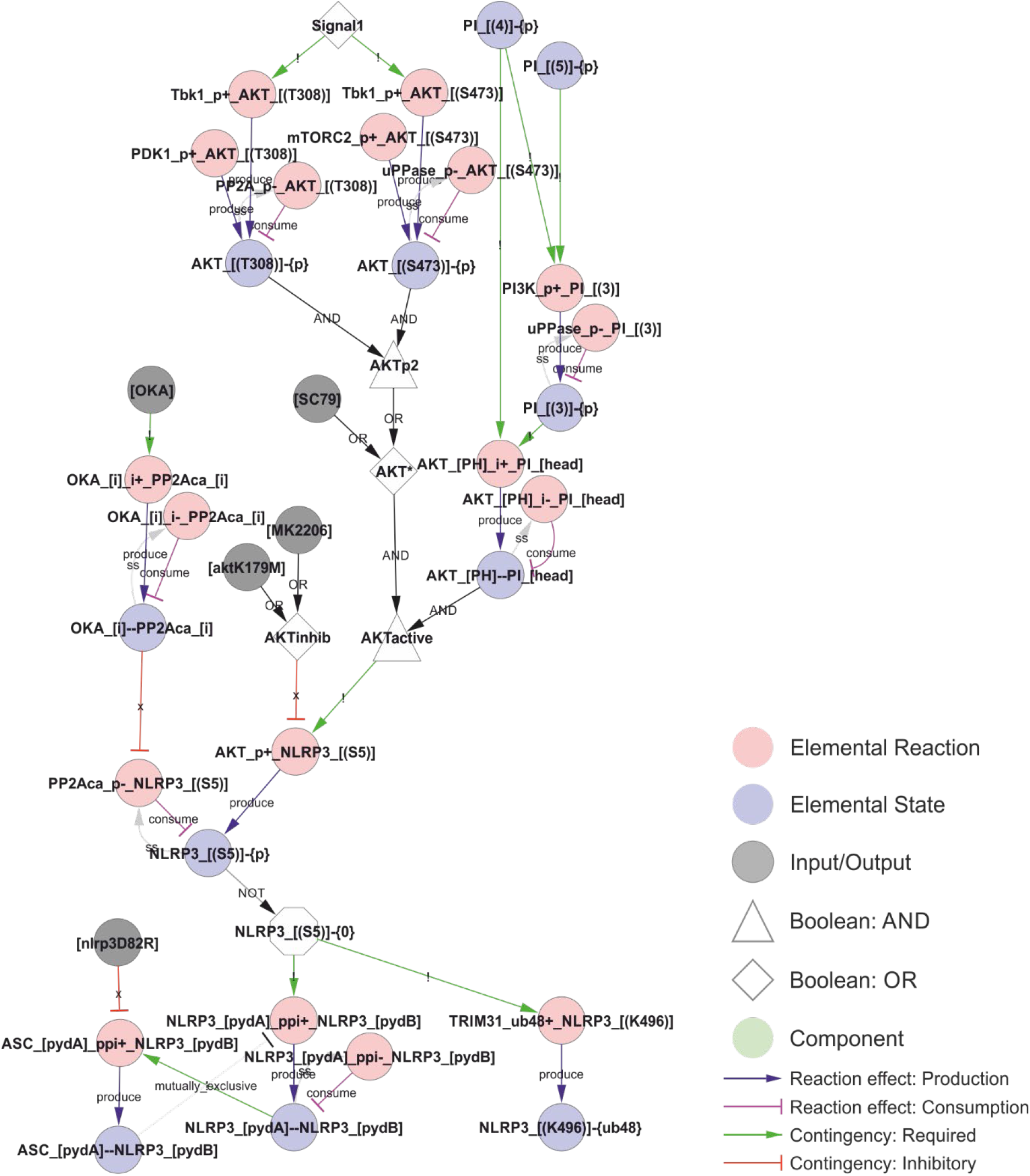
PYD polymerisation and ASC recruitment is inhibited by Ser5 phosphorylation. Inflammasome assembly requires formation of a NLRP3-PYD filament, which nucleates an ASC-PYD filament through homotypic PYD-PYD interactions, and these interactions are prevented by Ser5 phosphorylation. Ser5 phosphorylation is controlled by AKT, which is constitutively active at the plasma membrane. However, AKT activity is also increased by Signal 1-dependent Tbk1 phosphorylation, introducing a possible time delay to polymerisation upon priming. In addition, Ser5 phosphorylation has been reported to stabilise NLRP3 by preventing K48-linked polyubiquitylation of Lys496 and hence proteasomal degradation of NLRP3. *Walkthrough: Signal 1 (white node, top) activates Tbk1 to phosphorylate AKT, although this phosphorylation is also mediated (constitutively) by Pdk1 (at Thr308) and mTORC1 (at Ser473) and reversed by PP2A and an unknown phosphatase. Phosphorylation, together with binding to PI(3,4)P2 and PI(3,4,5)P3 (PI phosphorylated on position 3 AND 4), activates AKT (AKTactive, white node in the middle) to phosphorylate NLRP3 at Ser5. Only NLRP3 that are unphosphorylated at Ser5 (the white (NOT) octagon: “NLRP3_[(S5)]-{0}”) can support NLRP3-PYD polymerisation (through protein-protein interactions, “ppi+”, between the A and B sides of the PYD domain (pydA and pydB)) and K48 linked (“ub48+”) ubiquitylation at Lys496 of NLRP3 (“NLRP3_[(K496)]-{ub48}”). NLRP3 pyd polymers (represented by a single bond: “NLRP3_[pydA]-- NLRP3_[pydB]”) can then nucleate NLRP3-ASC PYD domain polymerisation: “ASC_[pydA]_ppi+_NLRP3_[pydB]”*.

In contrast to the inhibitory Ser5 phosphorylation, priming-dependent Ser198 phosphorylation is required for inflammasome assembly and IL-1β release (Figure 5). Ser198 is phosphorylated by JNK1 upon priming, and this phosphorylation is essential for self-association and hence oligomerisation of NLRP3 [19]. Phosphorylation of Ser198 also decreases ubiquitylation of NLRP3 [19], by inhibiting ubiquitylation, promoting deubiquitylation, or both. Hence, Ser198 phosphorylation may constitute a priming dependent switch between inflammasome assembly and degradation. The Ser198 residue is localised in the FISNA domain (residues 138-208) [81], which is the region that undergoes the largest structural changes during NLRP3 activation [44]. The fact that overexpression of NLRP3 can overcome this licensing requirement, strongly suggests that it is the deubiquitylation (negative licensing) rather than the phosphorylation (positive licensing) that control NLRP3 assembly. In the model, Ser198 phosphorylation is directly required for deubiquitylation of the leucin-rich repeats (LRR) domain of NLRP3 and, through this effect, indirectly promotes self-association of NLRP3.

**Figure 5:**
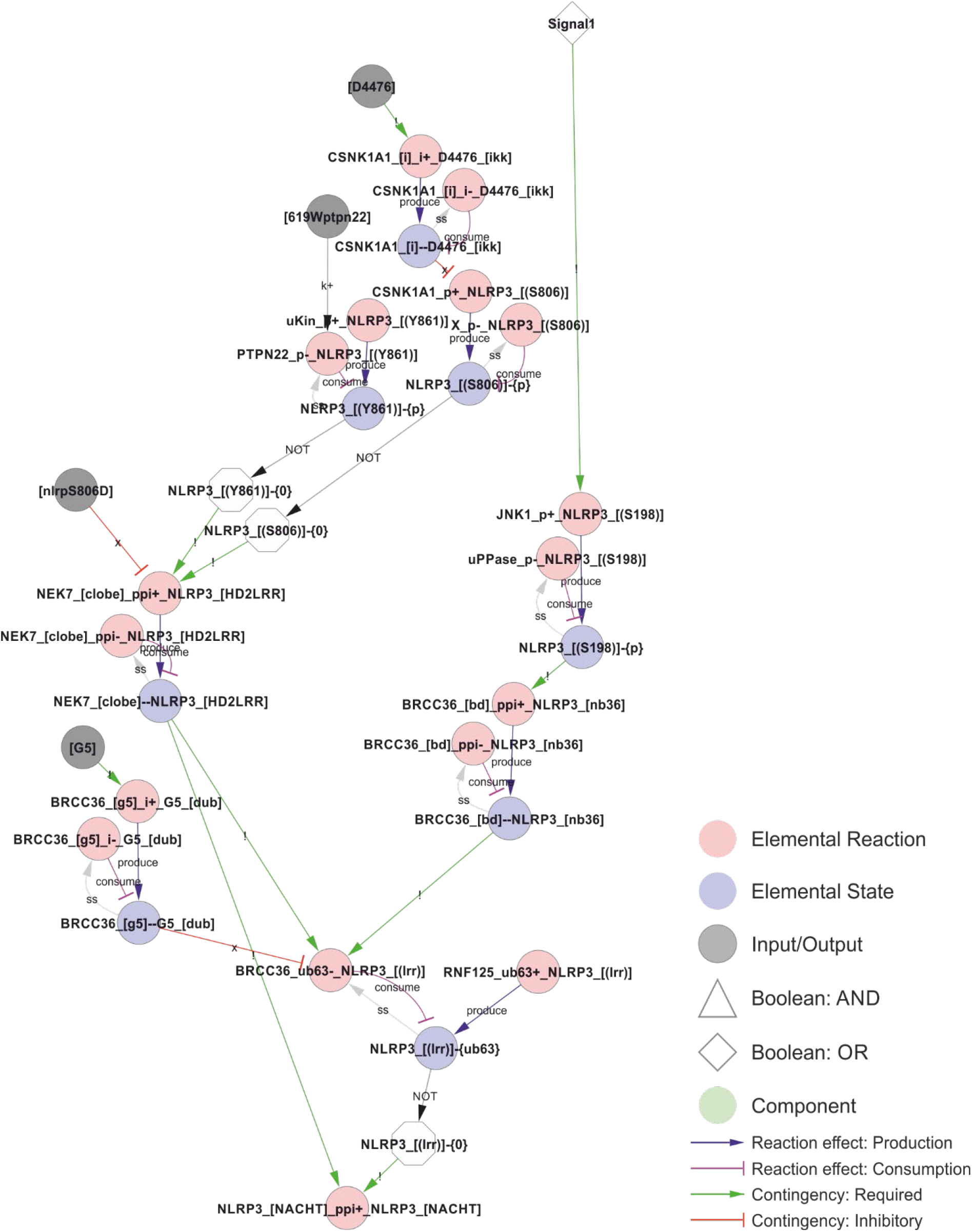
Control of NEK7 recruitment and BRCC36-dependent deubiquitylation of the LRR domain. Phosphorylation of Ser806 and Tyr861 prevents NEK7 recruitment and thereby BRCC36 dependent ubiquitylation, but does not appear to be regulated by Signal 1 or 2, suggesting that phosphorylation of Ser806 and/or Tyr861 may impose a spatiotemporal restriction on NLRP3 activation. In contrast, JNK1 dependent phosphorylation of S198 constitute a bona fide licensing event, as it depends on Signal 1 and is a prerequisite for BRCC36-dependent deubiquitylation of the LRR domain, for NLRP3 self-association (encoded as an indirect dependence in the model), and hence for assembly and activation of the inflammasome. *Walkthrough: Signal 1 (top right) triggers JNK1 phosphorylation of Ser198 to produce NLRP3_[(S198)]-{P}, which is required for BRCC36 bidning to NLRP3. The bond (“BRCC36_[bd]--NLRP3_[nb36]”) is in turn required for removal of the K63 linked polyubiquitin chains (“-{ub63}”) on the LRR domain of NLRP3 (“BRCC36_ub63-_NLRP3_[(lrr)]”), which consumes NLRP3_[(lrr)]-{ub63} and thereby produce the unubiquitylated form: NLRP3_[(lrr)]-{0} (white octagon; NOT of NLRP3_[(lrr)]-{ub63}) that is required for oligomerisation in the model. This deubiquitylation also requires that NLRP3 is bound to NEK7 (“NEK_[clobe]--NLRP3_[HD2LRR]”), and it is prevented by binding of the G5 inhibitor to BRCC36 (“BRCC36_[G5]—G5_[dub]”) which only occurs in the presence of G5 (input node: “[G5]”). Continuing upwards on the left side, NEK7 only binds to NLRP3 when the latter is unphosphorylated at residues Ser806 and Tyr861. Furthermore, the biding is prevented by the NLRP3S806D phosphomimic mutation (input: “[nlrp3S806D]”)*.

The Ser198-mediated effect converges with the effects of Ser806 and Tyr861 phosphorylation. Ser806 phosphorylation has been shown to prevent NEK7 interaction and, through this, to prevent BRCC36 dependent deubiquitylation of the LRR-domain of NLRP3 [22]. Again, ubiquitylation in NLRP3 prevents its self-association and hence the assembly of a signalling competent inflammasome. Ser806 is targeted by casein kinase (CSNK1a1), which is presumed to be constitutively activated [22], suggesting that phosphorylation of this site also may impose a spatiotemporal restriction on NLRP3 activation rather than constitute an actual priming event. Similarly, the Tyr861 phosphorylation interferes with NEK7 recruitment [21], but is not known to be regulated in response to either Signal 1 or 2 [59]. In the model, both Ser806 and Tyr861 must be unphosphorylated for NLRP3 to bind NEK7, and NLRP3 must bind NEK7 to allow BRCC36 mediated deubiquitylation of the LRR domain.

The phosphorylation of Ser295 has been shown to act as both a positive and negative regulator of NLRP3 activation (Figure 6). This is consistent with its role in controlling the ATPase activity of NLRP3, as both ATP binding and hydrolysis has been found to be important for NLRP3 activation [8, 68]. Ser295 is phosphorylated by both PKA [23] and PKD [16]. The Ser295 phosphorylation has been shown to prevent NLRP3 activation [23], but at the same time to be necessary for NLRP3 release from Golgi membranes and relocation to the compartment of activation, as discussed further below [16]. This implies that the ATPase activity or ADP-bound form helps NLRP3 to dissociate from the membrane, and that the Ser295 phosphorylation positively affects this. In the model, this is implemented as an inhibition by Ser295 phosphorylation of the ADP to ATP exchange, with the result that NLRP3 will favour the ADP bound form. Similarly, the NLRP3 inhibitor MCC950 is considered an ATPase inhibitor [70] but stabilises the inactive ADP bound form [82]. This is more consistent with an inhibition of ADP-to-ATP exchange, indirectly decreasing the ATPase activity through reduced exchange, and hence this is the mechanisms implemented in the model.

**Figure 6:**
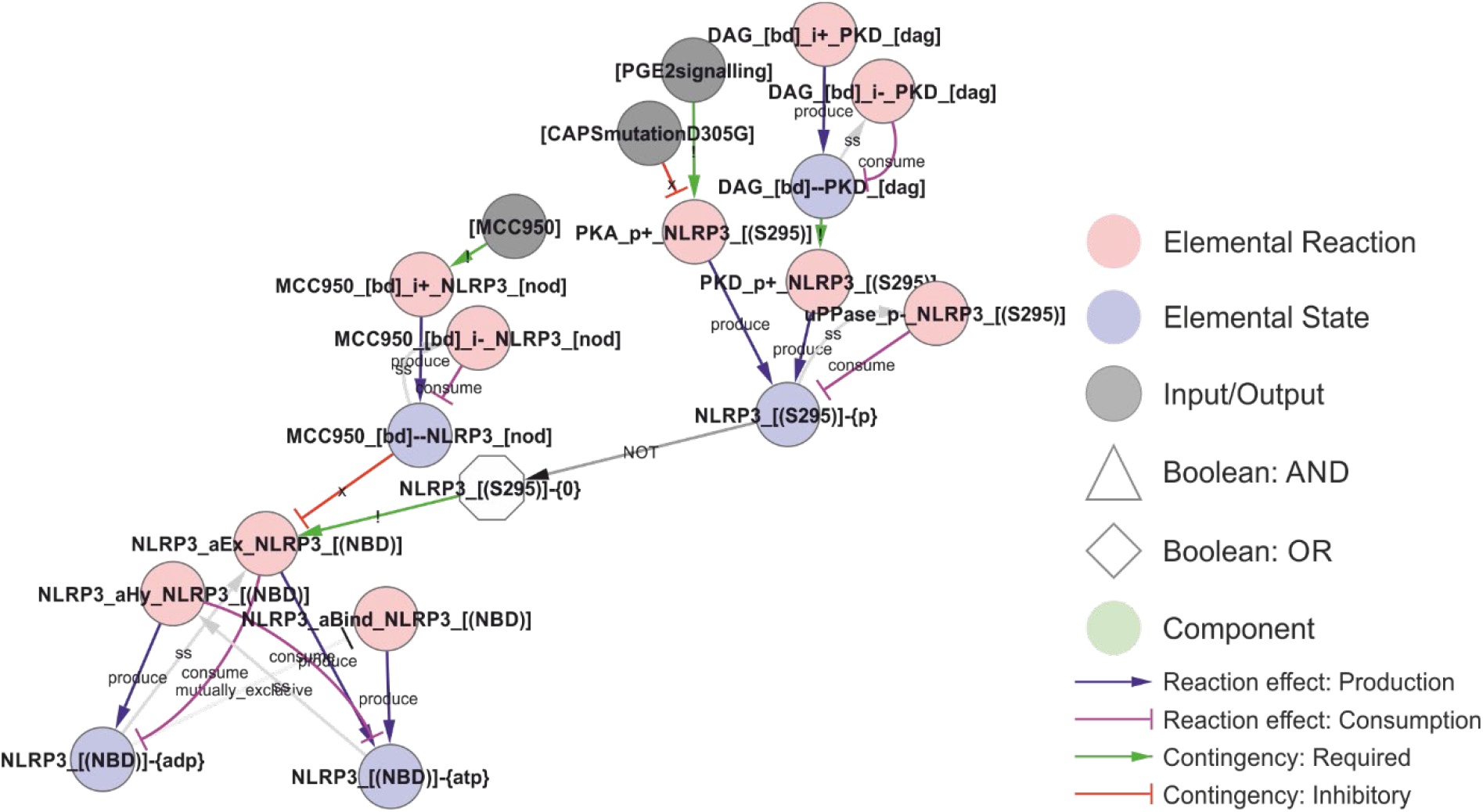
Regulation of the ATPase cycle by MCC950 and Ser295 phosphorylation. Phosphorylation of NLRP3 at Ser295 is controlled by both PKA and PKD, and it has been ascribed both positive and negative roles in NLRP3 regulation. These effects have been attributed to inhibition of the ATPase activity, but, like MCC950, it appears more likely that they stabilise the ADP-bound form by preventing nucleotide exchange – which would decrease the apparent ATPase activity without shifting NLRP3 towards an ATP-bound active form. *Walkthrough: The ATP-cycle of NLRP3 is represented by three reactions: (i) Binding of empty NLRP3 to ATP (“NLRP3_aBind_NLRP3_[(NBD)], (ii) APT hydrolysis (“NLPR3_aHy_NLRP3_[(NBD)]”), and ADP-to-ATP exchange (“NLRP3_aEx_NLRP3_[(NBD)]”). Note that nucleotide binding (-{0}/-{ATP}/-{ADP} = empty/ATP-bound/ADP-bound, respectively) is treated as an internal state of NLRP3, and that NLRP3 “acting on itself” appears both to the left and right in the reaction string. In the initial model, ADP-to-ATP exchange is inhibited by MCC950 binding (“MCC950_[bd]--NLRP3_[NOD]”) and requires that Ser295 is unphosphorylated (“NLRP3_[(S295)]-{0}"). Ser295 can be phosphorylated by both PKA and PKD. As described in the model analysis below, gap-filling suggested that also the ATP binding must be regulated to prevent activation of newly synthesised NLRP3*.

### The trigger signal: NLRP3 activation

After priming and licensing, NLRP3 can be activated by a wide range of triggers (“Signal 2”). It is an outstanding question if these triggers converge on one common signal, and, if so, what the actual trigger signal is. The known NLRP3 triggers can be divided into three general categories (Table 1), in this article labelled Type I - III: Type I triggers (e.g., nigericin or ATP via P2X7) cause ion fluxes, Type II triggers (e.g., MSU or LLOMe) causes lysosomal damage, Type III triggers (e.g., imiquimod and CL097) inhibit the mitochondrial electron transport chain. Some of these triggers can be suppressed by KCl, leading to the suggestion that K+ efflux constitutes the trigger signal, but some triggers – notably Type III triggers – are insensitive to external KCl [39]. Nigericin activation can also be blocked without affecting K+ efflux [123], showing that K+ efflux cannot be the actual trigger mechanism. Other proposed unifying mechanisms, such as ROS production, have also been discarded as exceptions have been discovered [109]. These findings suggest that either the three trigger classes activate NLRP3 in different ways, or, maybe more likely, that the trigger mechanism is even more fundamental.

The existence of such a fundamental feature of NLRP3 activation could be related to cellular energy metabolism. All NLRP3 triggers perturb cellular energy, either by increasing ATP consumption through uncoupling of transmembrane ion pumps (Types I and II) or by disrupting ATP production (Type III). In line with this, NLRP3 is an ATPase [68], ATP-binding is essential for activity [68], as is ATP hydrolysis [8]. The most potent NLRP3 specific inhibitor, MCC950, is thought to block NLRP3’s ATPase activity [70], although this is disputed [124]. In addition, specific phosphorylation at Ser295 blocks NLRP3’s ATPase activity [8], and several mutations in the vicinity of this phosphorylation site is found in Cryopyrin-associated periodic syndromes (CAPS) [23], which are associated with spontaneous NLRP3- dependent inflammation. The CAPS mutations in the nucleotide binding domain (NBD) appear to have a higher affinity for ATP and thereby to stabilise the open, ATP-bound conformation [125]. For example, the R262W mutation increases speck formation [124], consistent with the predicted increase in ATP-binding affinity [125]. However, most NLRP3 mutations that decrease the ATPase activity prevent speck formation [124]. As the ATP-bound form of NLRP3 has been established as active [44], these apparently contradictory findings suggest that also the ATPase cycle is important for NLRP3 function. However, as a chimeric NLRP3-NLRP6 protein, containing the NLRP6 NBD, responds to triggers in a similar manner as NLRP3 [18], it is unlikely that NLRP3 itself acts as an energy sensor. Instead, cellular energy may influence something else that NLRP3 is able to sense.

Approaching the problem from the other end, it is known that licensed NLRP3 can assemble and activate on two different lipids: cardiolipin (CL) [66] and phosphoinositol-4-phosphate (PI(4)P) [31]. Incidentally, these lipids are also targeted by gasdermin D [25], the pore forming executioner of pyroptosis and ultimate effector of NLRP3 inflammasome activation [30]. In the (casp8-/-) model presented here, where NLRP3 activation absolutely depends on NEK7 [78], we assume that NLRP3 would need to reach the centrosome, where NEK7 is reported to be localised [126], in order to be activated. Consistent with this hypothesis, NLRP3 activation has been found to depend on microtubule-based transport to bring NLRP3 to the centrosome and NEK7 [127]. It is worth stressing that caspase-8 is known to be recruited to and activated by CL [128], pointing to a possible mechanistic role of caspase-8 in NLRP3 activation as well as a potential difference of the NLRP3 activation by CL and PI(4)P. However, the mechanistic role of caspase-8 in NLRP3 activation remains uncertain [129], and there is evidence for a role for microtubule transport also of mitochondria in NLRP3 activation [130]. For the purpose of this model, NLRP3 needs to reach NEK7 at the centrosome for NLRP3 inflammasome activation both by PI(4)P and CL binding.

Cardiolipin (CL) is a bacterial lipid. In healthy eukaryotic cells, CL is only found in the inner mitochondrial membrane [131]. CL is only exposed to the cytoplasm and to NLRP3 binding after the outer mitochondrial membrane has been compromised, e.g. after the Mitochondrial membrane permeability transition (MPT) [132] and rupture of the outer mitochondrial membrane [133], or after CL translocation to the outer mitochondrial membrane in response to e.g. apoptotic stimuli [134]. Hence, in the absence of apoptotic signals, CL-mediated NLRP3 activation requires substantial mitochondrial damage or an intracellular bacterial infection. CL exposure in the mitochondrial membrane is a known damage signal inducing mitophagy [135], and in the context of infection (NLRP3 priming) it constitutes a danger signal. MPT may be triggered by a Ca^2+^-dependent pore opening, which is sensitised by e.g., oxidative stress or mitochondrial membrane depolarisation [133], offering an explanation to the conflicting reports on ROS regulation of NLRP3. Importantly, NLRP3 has been shown to interact with CL, and this interaction depend on an NLRP3 trigger signal (shown for type I (ATP) and II (Silica) triggers; [66], supporting the notion that NLRP3 triggers cause mitochondrial damage.

In contrast to CL, PI(4)P is constitutively present at cytoplasmic membranes and hence constitutively available for NLRP3 binding. Under normal conditions, it is found associated with the plasma membrane and Golgi [136], and it accumulates in autophagosomes [137], Rab7 positive late endosomes/lysosomes [136], and in Rab7 positive late-stage phagosomes [138]. Exposure of cells to NLRP3 triggers leads to an accumulation of PI(4)P-containing vesicles, and this accumulation is independent of NLRP3 [31]. These vesicles were initially thought to represent a dispersed trans-Golgi network (dTGN) due to the presence of the TGN38/46 marker, but TGN38/46 itself shuttles to the plasma membrane and back through endosomes, and this transport is known to be impaired by NLRP3 triggers [139]. Hence, these compartments likely correspond to PI(4)P-containing late endosomal Rab7 positive vesicles [136], and they likely accumulate due to endosomal trafficking defects [140]. NLRP3 triggers are known to impair endosomal acidification [39], and impaired endosomal acidification has been shown to impair trafficking and to prevent return of TGN38/46 to the TGN [141], explaining the accumulation of these vesicles. Even in plants, V-ATPase inhibition leads to accumulation of vesicles [142] that are enriched in PI(4)P [143], suggesting that this is a highly conserved and hence fundamental process. Consistently, genetic disruptions of endosomal trafficking, resulting in accumulation of PI(4)P containing endosomes, trigger the NLRP3 inflammasome upon priming [34]. Importantly, the assembly of NLRP3 on late endosomal vesicles enables microtubule-based transport to the centrosome [144], possibly via Rab7/RILP interaction with Dynein [145], where NLRP3 can interact with NEK7 to assemble an active inflammasome [60]. This mobility may be the key difference to NLRP3 recruitment to PI(4)P on the plasma membrane or TGN. In line with this, permanent PI(4)P localisation did not activate NLRP3 without a trigger [31]. In fact, release of NLRP3 from Golgi PI(4)P was essential for activation [16]. Importantly, the accumulation of TGN38/46 positive endosomal vesicles has been observed for all three types of NLRP3 triggers: (I) nigericin, (II) LLOMe, and (III) imiquimod, making it the most general feature of NLRP3 activation yet identified [140]. However, it leaves the question to why those vesicles accumulate.

Again, the evidence points towards cellular energy: i) endolysosomal acidification requires ATP, ii) intracellular ATP has been shown to decease in response to NLRP3 stimuli through K^+^- and Ca^2+^- mediated mitochondrial dysfunction, and iii) artificial decrease of intracellular ATP through inhibition of glycolysis has been shown to trigger NLRP3 [146]. Moreover, triggering of NLRP3 through P2X7 involves mobilisation of mitochondrial potassium [17]. Recently, ATP-generation in mitochondria was found to be driven to a large extent by the secondary K^+^ gradient [37] (generated by the mitochondrial H^+^/K^+^ antiporter [54]), providing a possible mechanistic explanation to how potassium outflux could lead to a strong and immediate reduction of ATP production. This also provides a possible explanation for why Type III triggers are insensitive to KCl supplementation, as inhibition of the mitochondrial electron transport chain (ETC) would disrupt both the primary H^+^ and the secondary K^+^ gradient over the mitochondrial inner membrane. In addition, NLRP3 triggers cause NAD^+^ to decrease, and NAD^+^ supplementation has been shown to prevent NLRP3 activation [130]. This effect has been found to depend on SIRT2 and possibly on α-tubulin deacetylation, suggesting that low NAD+ drives dynein dependent transport towards the centrosome [130]. However, NAD^+^ is also an essential cofactor in glycolysis and a drop in reoxidation of NADH through ETC inhibitors (like the NLRP3 triggers Imiquimod and CL097 [39]) would lead to a strong reduction in glycolytic ATP production as well. It is worth pointing out that endolysosomal pumps appear to be supplied at least partially by endolysosomally localised GAPDH, which uses NAD^+^ as a cofactor to produce ATP [147]. Moreover, GAPDH inhibition can – as inhibition of glycolysis in general – trigger NLRP3 activation [146, 148], although this of course have an impact on global energy levels. Consistently, direct inhibition of the lysosomal V-ATPase has been shown to trigger NLRP3, and this activation cannot be prevented by external supplementation of K^+^ [149]. This would be consistent with a model in which decreasing cellular energy levels leads to loss of (internal, proton-gradient driven) membrane potential, which in turn is sensed through its effect on intracellular trafficking. Hence, the above evidence converges on a scenario where NLRP3 responds to energy by detecting compromised membranes that are rerouted due to the lack of appropriate membrane potential.

The evidence on a role for cellular energy in inflammasome activation is also consistent with disruption of another key physiological feature: osmotic integrity. Animal cells lack cell walls, and hence must maintain isoosmolarity at all times or they will shrink, swell, or rupture [150]. As cells contain large amounts of osmotically active compounds, such as proteins and metabolites, they must compensate these with a net gradient of inorganic ions to maintain isoosmolarity [151]. The primary architect of this gradient is the Na^+^/K^+^-ATPase that establish a gradient over the plasma membrane. The relatively high permeability to K^+^ allows this ion to approach its equilibrium potential, maintaining an electric gradient over the plasma membrane. This in turn drives the outflow of Cl^-^, and this creates the “osmotic room”, which is necessary for accommodating all essential biomolecules within the cytoplasm while maintaining a zero osmotic pressure over the plasma membrane [152]. If this gradient is compromised, the cell will start taking up water, swell, and eventually burst, and, consequently, detection of this gradient collapse is an acute danger signal. Several lines of evidence suggest that such a gradient collapse could be the fundamental trigger for NLRP3 activation. First, NLRP3 is activated by hypotonicity, and this activation is conserved from mammals to fish [123]. Second, hypertonic solutions prevent NLRP3-dependent IL-1β release in response to triggers such as nigericin and MSU [123]. Third, the “symptoms” of hypoosmotic stress is similar to NLRP3 activation, including release of KCl in order to reduce the volume of “osmotically swollen” cells [151]. Fourth, the endolysosomal system, the site of (PI(4)P dependent) NLRP3 activation, is considered the “osmometers” of the cell, playing a key role in adaptation to and survival in hypotonicity [153]. Fifth, Rab7 positive vesicles are involved in the acute response to hypotonicity, by absorbing water and excreting it on the outside, and this function relies on the V-ATPase as inhibition with Baf-A1 inhibits vacuolisation and massively increase cell death in response to hypoosmotic shocks [154], mirroring the conditions of NLRP3 activation. Hence, the activation of NLRP3 on PI(4)P positive endolysosomes could indicate an osmotic problem.

This reasoning leaves the question to how the NLRP3 triggers affect the inorganic ion gradient. The simplest answer may again be cellular energy. The maintenance of the inorganic ion gradient over the plasma membrane is energy expensive (estimated 25-75% of the total cellular ATP consumption, depending on cell type), and the pump activity is known to strongly depend on the intracellular ATP concentration [155]. A drop in Na^+^/K^+^-ATPase activity below the threshold needed to maintain this Cl^-^ gradient will invariably lead to an influx of water, cell swelling, and, eventually, cell rupture [150]. Indeed, the decreased activity of the Na^+^/K^+^-ATPase has been proposed as the main mechanism mediating ATP deficiency-induced apoptosis [156]. Consistently, inhibitors of the Na^+^/K^+^-ATPase, such as Ouabain or Digoxin, trigger NLRP3 after priming with LPS [157]. Importantly, significant K^+^ efflux requires a collapse of the electrical gradient or compensatory transport of other ions, and hence ATP/P2X7 (cation influx) dependent IL-1β activation does not occur in the absence of extracellular Na^+^ and Ca^2+^ [40]. Along the same lines, depletion of extracellular Cl^-^ and K^+^, but neither alone, suffice to activate NLRP3 in LPS primed cells [158]. Indeed, different Cl^-^-channel inhibitors can at least diminish NLRP3 activation in response to nigericin [159], ATP [160], and MSU [160], highlighting the importance of co-or counter-fluxes for K^+^ efflux. It is worth to mention that disruption of the normal regulatory volume decrease (RVD), through knock-out of the volume-regulated anion channel (VRAC [153]) LRRC8A, prevents NLRP3 activation by hypoosmotic stress but has no effect on activation by type I, II or III stimuli [159], which would be consistent with an insufficient pump activity as the underlying cause in the latter cases. Furthermore, deletion of WNK1, a Cl^-^-responsive regulator of Cl^-^-cation-cotransporters that balance intracellular Cl^-^ and K^+^, aggravates NLRP3 activation in response to type I (ATP, nigericin), type II (MSU) and type III (imiquimod) triggers [161]. This shows that Cl^-^ fluxes play an important role in NLRP3 activation. However, at least one Cl^-^-channel inhibitor (DIDS) can block NLRP3 activation without preventing loss of intracellular Cl^-^, suggesting that the ion fluxes over internal membranes is the critical determinant for NLRP3 activation [160]. In fact, type II stimuli has been shown to induce K^+^ leakage into the lysosomes, causing hyperpolarisation and lysosomal rupture [32] that precedes NLRP3 activation [162]. Of note, the lysosomal integrity can be rescued by increasing internal K^+^ through high external KCl [32]. Furthermore, phosphatidylinositol-4 kinase type 2a (PI4K2A) accumulates rapidly on damaged lysosomes after LMP, to generate high levels of PI(4)P [163] that allow the compromised lysosome to serve as a platform for NLRP3 activation. Hence, we hypothesise that NLRP3 is activated by intracellular trafficking problems originating from compromised osmotic regulation of the intracellular membranes, which are tagged by PI(4)P.

How does this then lead to NLRP3 activation? A recent study found that the N-terminal part of NLRP3 was sufficient to impose nigericin (type I), MSU (type II), and imiquimod (type III) regulation of a NLRP3- NLRP6 chimera [18]. Inclusion of residues 1-132 of NLRP3 is sufficient for at least partial regulation that is sensitive to external KCl for type I and II, but not type III, stimuli. Conversely, deletion of residues 92- 148, but not residues 92-132, is sufficient to completely abolish this regulation in NLRP3 [18]. This region of NLRP3 contains one notable feature: the KMKK^132^ (mouse KKKK) motif required to recruit NLRP3 to PI(4)P [31]. This motif, however, does not seem critical in human cells, which may be due to a second polybasic region (RKKYRKYVRSR^145^; [18]), and this redundancy would explain the apparently contradictory observation that residues 1-132 is sufficient but not required for regulation. Both these motifs lie within the short stretch required and sufficient to impose NLRP3-like regulation on the NLRP3-NLRP6 chimera, i.e. to make the chimera responsive to LPS plus nigericin in a KCl dependent manner [18]. Hence, these data strongly suggests that the common trigger of NLRP3 is its recruitment to PI(4)P. However, this recruitment is required, but not sufficient, for inflammasome activation [31], likely reflecting the need for assembly on mobile vesicles that can reach the centrosome and the essential interaction partner NEK7. Based on this reasoning, we propose a model where membrane recruitment and transport to the centrosome combine into a trigger signal for NLRP3 (Figure 7).

**Figure 7:**
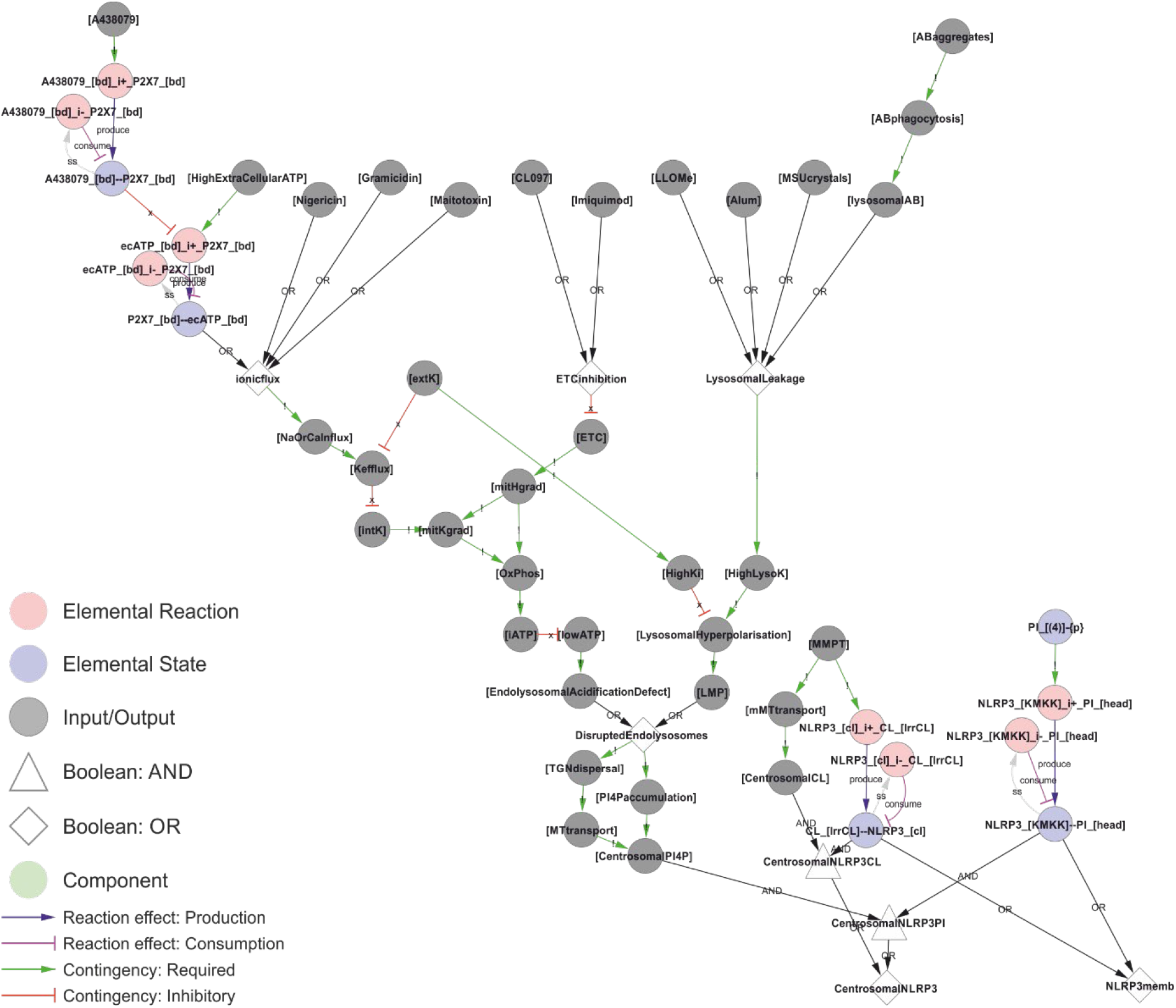
A unifying hypothesis for NLRP3 inflammasome activation. Based on current molecular knowledge, we propose that the common feature of all NLRP3 triggers is that they cause osmotic disruption of internal membranes and transport of those compartments to the centrosome for interaction with NEK7. For cardiolipin (CL)-dependent recruitment (right), this scenario would require mitochondrial membrane permeabilization (MMPT) to expose CL for interaction with NLRP3 and mitochondrial transport to the centrosome and NEK7. However, this is highly tentative as discussed below. For phosphoinositol-4-phosphate (PI(4)P)-dependent recruitment, the evidence is stronger as discussed in the text. Briefly: Type I triggers cause Na^+^ and/or Ca^2+^ influx, which in turn allow K^+^ efflux. Sinking intracellular potassium levels (which can be suppressed by external KCl) impair ATP generation in the mitochondria, and the sinking energy levels – possibly in combination with uncontrolled ion fluxes over internal membranes – disrupts endolysosomal acidification and hence osmotic control. Type III triggers achieve the same outcome by direct inhibition of the ETC. Type II triggers have been shown to directly destabilise lysosomes, which burst due to the rising osmotic pressure. Type III triggers could also disrupt the mitochondrial osmotic control, triggering MPT and CL exposure, together with mitochondrial transport to the centrosome upon failed mitophagy. Similarly, ionophores could directly destabilise internal compartments such as endosomes with the same result. In all these scenarios, the ultimate reason for NLRP3 activation would be osmotic lysis or loss of membrane potential, resulting in an accumulation of compromised internal compartments that accumulate PI(4)P and are mobile enough to reach the centrosome for inflammasome activation. *Walkthrough: At the top of this figure, we have the trigger signals, and at the bottom the two outputs that need to occur simultaneously to active NLRP3 in the model: Membrane recruitment of NLRP3 (“NLRP3memb”), and transport of that membrane to the centrosome (“centrosomalNRLP3”). Type I triggers converge on “ionicflux” (an OR-gate; either trigger suffices), causing Na+ or Ca2+ influx, which in turn causes K+ efflux unless prevented by high levels of external K+ (“extK”). Loss of internal K+ (“intK") leads to an impaired mithochondrial K+ gradient (“mitKgrad”), impaired oxidative phosphorylation and lower cellular ATP (“iATP”). Energy production can also be impaired directly by inhibition of the electron transport chain by type III triggers (“ETCinhibition”), which directly impair the primary H+ gradient in the mitochondria (“mitHgrad”). Finally, type II triggers cause lysosomal leakage, leading to increasing levels of lysosomal K+ (“HighLysoK”) that, unless intracellular K+ is also high (“HighK”), leads to lysosomal hyperpolarisation and permeabilisation (“LMP”). Low ATP leads to an endolysosomal acidification defect that, like LMP, leads to disrupted endolysosomes that disperse (“TGNdispersal”) and accumulate PI(4)P (“PI4Paccumulation”), the dispersed endolysosomes can be transported to the centrosome, allowing for centrosomally localised PI(4)P: “CentrosomalPI4P”. This, together with NLRP3 binding to PI4P (“NLRP3_[KMKK]--PI_[head]”) constitutes the trigger mechanism in the model. A parallel mechanism is implemented for CL-based NLRP3 recruitment, but this is – as discussed below – highly tentative*.

### NLRP3 inflammasome activation and assembly

The above suggested NLRP3 activation process points to three key aspects of inflammasome activation: lipid binding, self-association, and assuming or stabilising the (ATP-bound) open conformation. PI(4)P binding allows NLRP3 to assume an open conformation (as measured e.g. by BRET [18]), which exposes its nucleotide binding domain and in turn allows ATP binding. ATP stabilised the open conformation as it is incompatible with the closed structure [44]. Hence, ADP hydrolysis is required for closing, and the intrinsic ATPase activity will return the monomeric NLRP3 to its closed state. We hypothesise that the combination of NEK7 and PI(4)P binding suffices to stabilise the open conformation and, in the context of Ser198 phosphorylation and LRR deubiquitylation, allows the formation of structured higher-order NLRP3 complexes. The importance of the ATPase cycle could be explained if ATP hydrolysis (i) is required for dissociation from PI(4)P and hence to sample different membranes, or (ii) shifts the affinity towards (de)ubiquitinating enzymes and hence the balance between signalling and degradation, or (iii) both. There is indeed evidence for both of these scenarios: phosphorylation of Ser295, which seems to favour ATP hydrolysis, has been shown to be essential for release of NLRP3 from Golgi membranes – where NLRP3 is not activated – to allow assembly of functional inflammasomes elsewhere [16]. Conversely, Ser295 phosphorylation inhibits activation, likely by stabilising the ADP-bound conformation as it is linked both to NLRP3 inhibition and decreased ATP turnover [23]. At the same time, the ATP cycle is associated with the transitions between open and close conformations. It is easy to envisage that deubiquitylation by BRCC36, which binds to the NACHT domain and acts on the LRR domain [61] is limited to the closed conformation. Both these hypotheses are implemented in the model (Figure 8).

**Figure 8:**
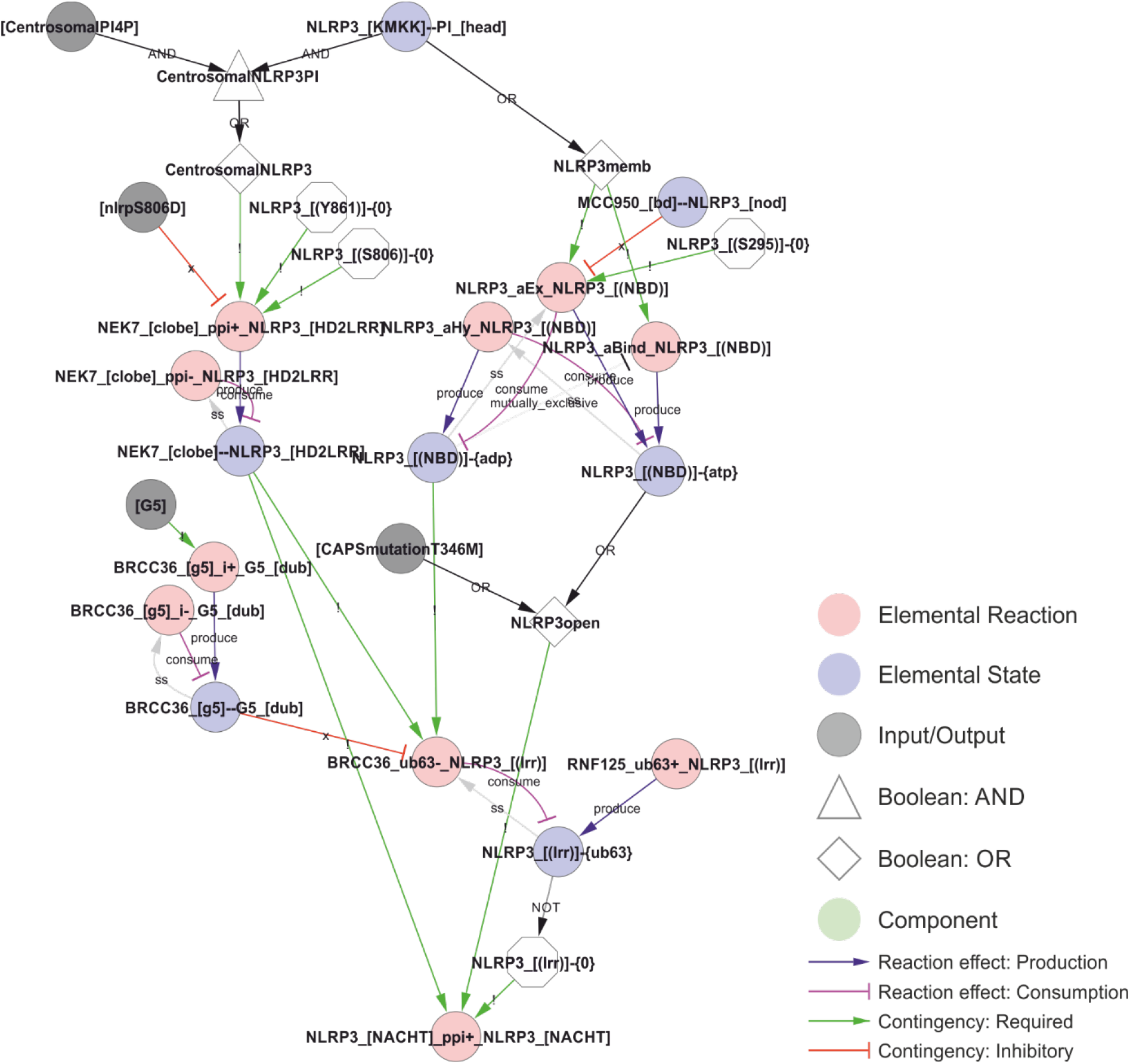
A model of the activation of NLRP3. The combination of phosphoinositol-4-phosphate (PI(4)P)-accumulation on and transport of compromised vesicles to the centrosome (“centrosomalPI4P”) with NLRP3 recruitment to PI(4)P localises NLRP3 to the centrosome where it can interact with NEK7. This interaction is controlled (prevented) by phosphorylation of NLRP3 in Ser806 or Tyr861, potentially restricting inflammasome activation in space, time, or intensity. At the same time, PI(4)P binding stimulate the transition of NLRP3 to the open conformation and the exchange of ADP to ATP. This transition and/or exchange is prevented by MCC950 or phosphorylation on Ser295, which stabilises the closed/ADP-bound state. The model also includes the effect of the ATP cycle on NLRP3 translocation as a requirement of NLRP3 dissociation from PI(4)P (and hence membranes) on ATP hydrolysis (the ADP bound state in the model) and by limiting deubiquitylation to the closed (ADP bound) conformation. Ser198 phosphorylation (Signal 1 priming via JNK1) is modelled as absolutely required – in combination with NEK7 binding – to allow BRCC36-mediated deubiquitylation of NLRP3s LRR domain, which in turn is necessary for self-association of NLRP3 – the first step of inflammasome assembly in the model. *Walkthrough: NLRP3 binding to head group of PI at the polybasic KMKK domain (“NLRP3_[KMKK]--PI_[head]”) leads to NLRP3 membrane recruitment (“NLRP3memb”) and, together with centrosomal PI4P localisation, to centrosomal NLRP3 (“CentrosomalNLRP3”). On the left side; centrosomal NLRP3 allows interaction with NEK7 (“NEK7_[clobe]_ppi+_NLRP3_[HD2LRR]”) dependent on the phosphorylation status of NLRP3’s LRR domain (see* Figure 5*). On the right side; NLRP3 membrane recruitment is – in the model – required for ATP binding and ADP-to-ATP exchange. ATP binding (“NLRP3_[(NBD)]-{ATP}”) in turn stabilises the open conformation (“NLRP3open”), which can also be achieved through the CAPS mutation NLRP3T346M). The open conformation in combination with NEK7 stimulates NLRP3 oligomerisation (“NLRP3_[NACHT]_ppi+_NLRP3_[NACHT]”), when the K63-linked ubiquitin chains are removed from the LRR-domain (“NLRP3_[(lrr)]-{0}”). This (“BRCC36_ub63-_NLRP3_[(lrr)]”) only occurs in the ADP bound (closed) conformation. Hence, the requirement for the ADP/ATP cycle is encoded through the requirement for the different states (ADP-or ATP-bound) at different stages of the activation process*.

Inflammasome assembly starts with the self-association of NLRP3 molecules, which, in the open and ATP-bound conformation, leads to the polymerisation of the NLRP3 PYD domains in a helical structure of about six PYD monomers per turn [29], implying that more than six NLRP3 molecules are needed to form a PYD filament. This initial PYD filament can then recruit ASC PYD domains to nucleate an ASC PYD filament [24]. The elongation is unidirectional, as ASC only elongates from the B-end of the NLRP3 PYD helix [29]. Hence, these filaments have a polarity. Furthermore, the formation is irreversible, as they do not disassemble upon dilution [29]. The proximity of the ASC monomers allows the ASC CARD domains to associate and create platforms for caspase-1 CARD filaments, bringing the caspase domains into proximity for dimerization, trans-autocleavage, and activation [24]. The model implementation of NLRP3 inflammasome assembly is shown in Figure 9.

**Figure 9:**
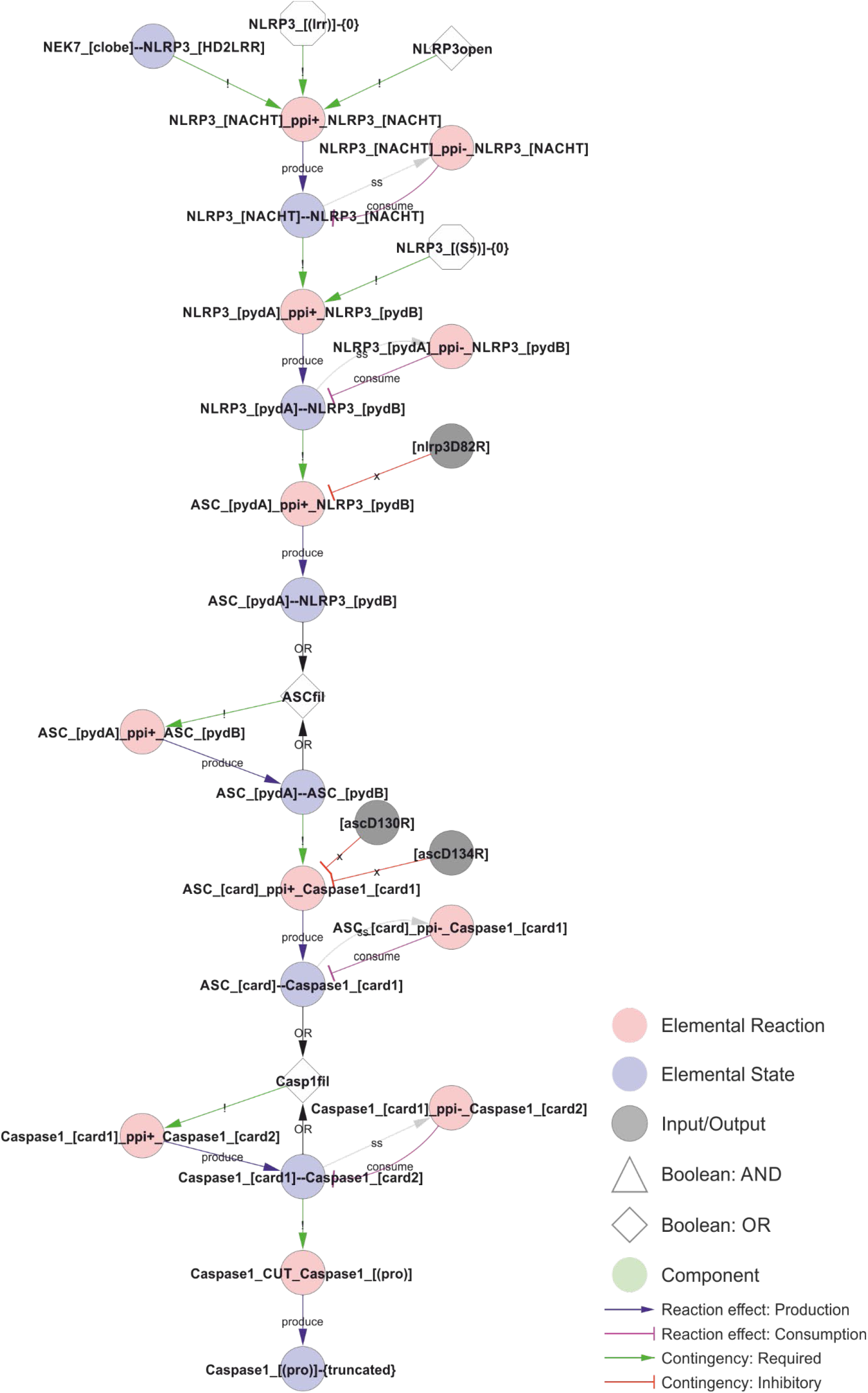
Model of the NLRP3 inflammasome assembly. The initial interaction between adjacent NLRP3ś NACHT domains requires an open conformation, NEK7 binding, and a lack of LRR domain ubiquitylation. The open conformation normally corresponds to the ATP-bound form, but is likely mimicked by certain CAPS mutations, such as R262W or T346M, allowing activation of NLRP3 after priming by LPS alone. The stable core NLRP3 interaction allow polymerisation of the NLRP3 PYD domains in the absence of Ser5 phosphorylation, which in turn nucleate the ASC PYD domain filament. ASC-ASC proximity provide the foundation for CARD filaments and nucleate caspase-1 polymerisation, which in turn allow caspase-1 cleavage and activation. *Walkthrough: The combination of NEK binding, open conformation, and the lack of K63 linked ubiquitin in the LRR-domain allows NLRP3 oligomerisation. The dimerisation of the NACHT domain (“NLRP3_[NACHT]--NLRP3_[NACHT]”) allows the polymersiation of the PYD domains, and the internal PYD polymers (“NLRP3_[pydA]-- NLRP3_[pydB]” nucleates the ASC-filament through ASC-NLRP3 (“ASC_[pydA]--NLRP3_[pydB]") and eventually ASC-ASC (“ASC_[pydA]--ASC_[pydB]”) interactions. The ASC polymers in turn nucleates caspase-1 filaments, first through ASC-caspase-1 (“ASC_[card]--Caspase1_[card1]”) and then caspase-1-caspase-1 (“Caspase1_[card1]--Caspase1_[card2]”) CARD-CARD interactions. Finally, the caspase-1 filaments trigger proximity-induced autocatalytic processing of caspase-1 (“Caspase1_cut_Caspase1_[(pro)]”) resulting in the truncated (“Caspase1_[(pro)]-{truncated}”) and active form of caspase-1*.

### NLRP3 inflammasome effectors and output

The output of NLRP3 signalling is mediated by three effectors that are activated by caspase-1 through proteolytic cleavage: IL-1β, IL-18, and gasdermin D (Figure 10; reviewed in [164]). Pro-IL-1β and pro-IL-18 are cleaved into their active, signalling competent form. They are leaderless and released into the extracellular space via exocytosis, plasma membrane pores, and/or cell rupture, to exercise a local or systemic pro-inflammatory effect [165]. In contrast, gasdermin D exerts its effect locally: the N-terminal peptide inserts in target membranes to form pores that are large enough to release IL-1β and IL-18 [30, 166], and to allow uncontrolled ion fluxes, in a similarly manner as type I or type II triggers. Of note, gasdermin D preferentially targets PI(4)P, PI(4,5)P_2_, and CL and also, but with apparently weaker affinity, phosphatidic acid and phosphatidylserine [25]. The overlap in lipid affinity with NLRP3 is striking, and this, in combination with the reported localisation of gasdermin D to the NLRP3 inflammasome complex [67], suggests that gasdermin D might be directed to target the membranes recruiting NLRP3. Hence, the NLRP3-gasdermin D axis could constitute an intracellular defence system designed to kill and dispose of – through autophagy – intracellular pathogens. Consistently, gasdermin D is cytotoxic and has the ability to kill bacteria [30]. It is also worth pointing out that gasdermin D pores allow ion fluxes that lead to further inflammasome activation in a positive feedback loop, unless both the original insult and signal are disposed of through autophagy. Consequently, caspase-4 and caspase-5, which can activate gasdermin D in response to intracellular LPS, indirectly triggers NLRP3 activation through gasdermin D and the ion fluxes it causes [167] (Figure 11).

**Figure 10:**
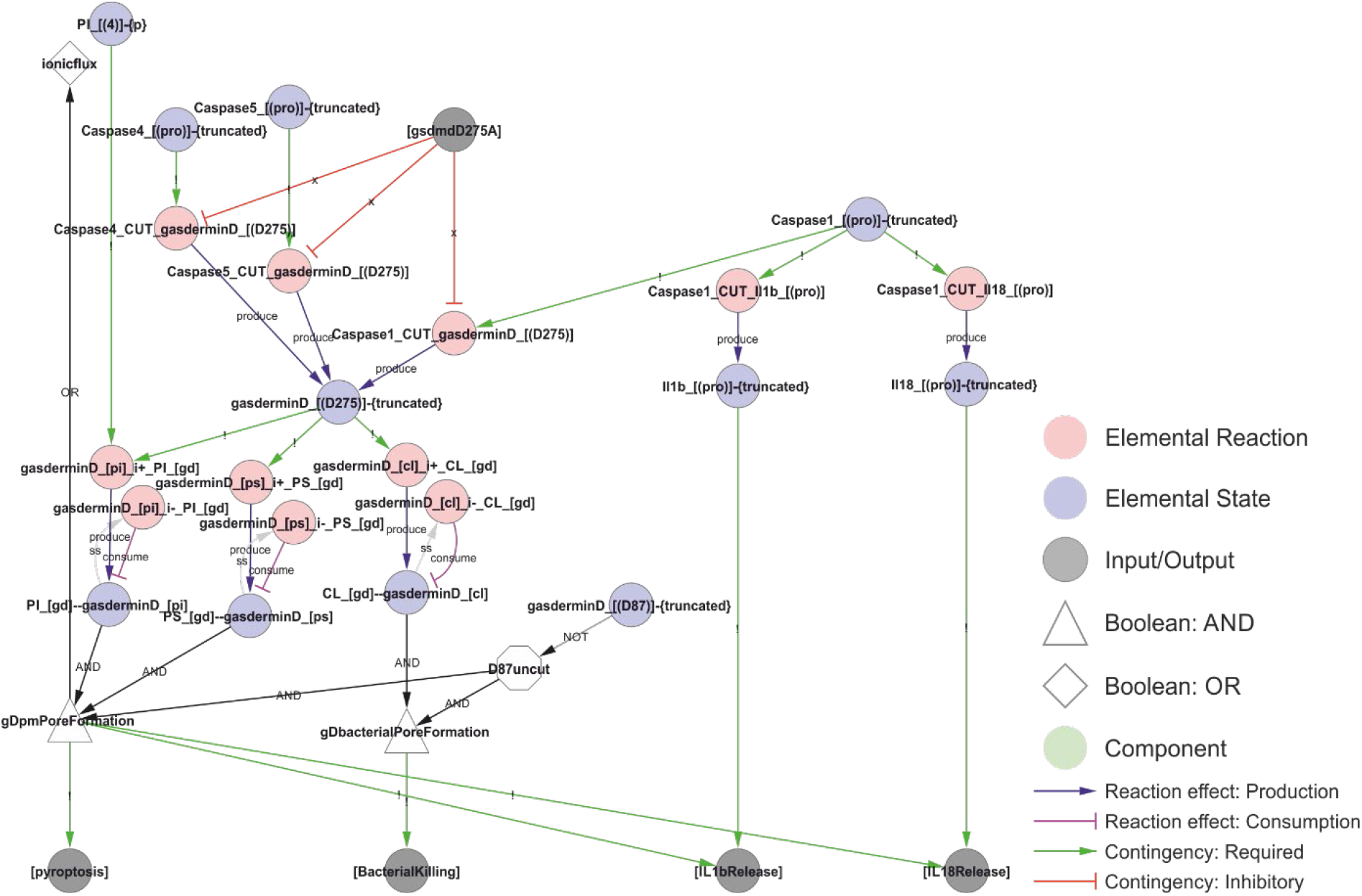
A model of NLRP3 output. Activated caspase-1 cleaves gasdermin D, pro-IL-1β, and pro-IL-18. Gasdermin D insertion in bacterial membranes can directly kill bacteria. Insertion in internal membranes may allow access to pathogens but may also kill the cell through pyroptosis. Cell lysis or insertion of gasdermin D pores in the plasma membrane will enable the release of IL-1β and IL-18 to the extracellular space. Formation of gasdermin D pores will allow ion flux that act as an NLRP3 trigger, creating a positive feedback loop. *Walkthrough: Truncation of Caspase-1, 4 and 5 (“Caspase1_[(pro)]-{truncated}”, etc.) is required to cut gasdermin D (“Caspase1_cut_gasderminD_[(D275)]”, etc.). The truncated form of gasdermin D binds PI, PS and CL. CL binding attacks intracellular bacteria (“BacterialKilling”). PI and PS binding results in internal and eventually plamsa membrane pore formation (“gDpmPoreFormation”), triggering the output of pyroptosis, IL-1β release and IL-18 release. Furthermore, it leads to ion fluxes (a Type I trigger), causing a positive feedback loop and explaining why intracellular LPS and Capsase-4/5 can activate the NLRP3 inflammasome (see* Figure 11*). Finally, IL-1β and IL-18 release (outputs) require that the interleukins have been cut by Caspase-1 (“Il1b_[(pro)]-{truncated}”)*.

**Figure 11:**
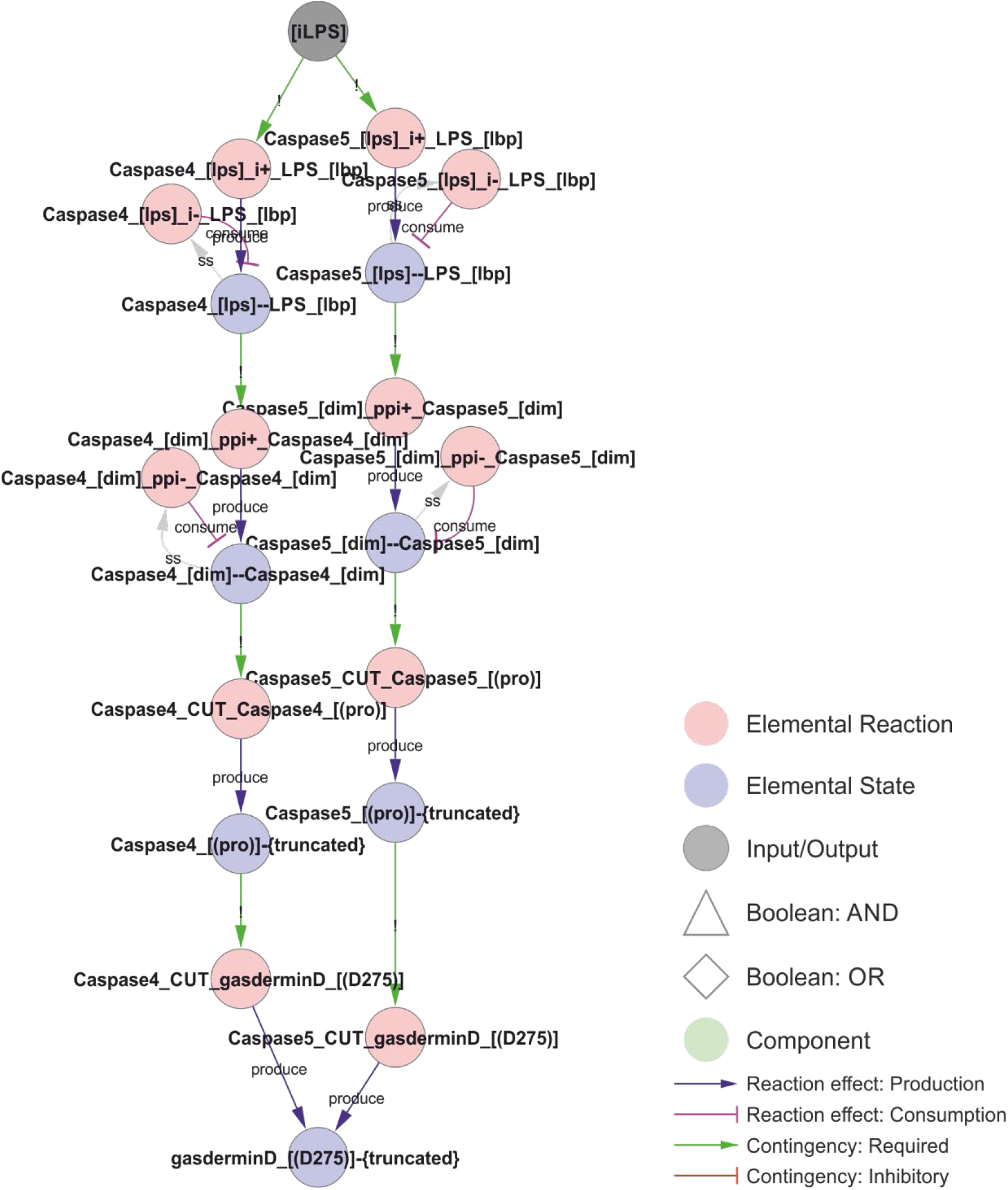
Non-canonical NLRP3 activation. Intracellular LPS can bind to and activate caspase-4 or caspase-5. LPS induces caspase dimerization, trans-autocleavage, and activation. Activated caspase-4 or 5 cuts and activates gasdermin D, which inserts into the membrane causing ionic fluxes and hence constitutes a canonical NLRP3 trigger. *Walkthrough: LPS binding to Caspase-4 (“Caspase4_[lps]-- LPS_[lbp]”; left) or Caspase-5 (right) triggers Caspase homodimerisation (“Caspase4_[dim]-- Caspase4_[dim]”, which in turn leads to proximity-induced cleavage and activation, as the truncated forms of Caspase-4 and 5 (“Caspase4_[(pro)]-{truncated}”) can cut and activate gasdermin D*.

### NLRP3 turnover

NLRP3 degradation seems to occur through both proteasomal and autophagosomal degradation, and to be controlled at several levels including through ubiquitylation (Figure 12). NLRP3 has also been reported to be targeted for precision autophagy in a ubiquitin independent manner via interaction with TRIM20 (pyrin) [26, 62]. As TRIM20 mutations are associated with familial Mediterranean fever [168], and as TRIM20 interacts with both ASC and NLRP3 [62] and links those to the autophagy machinery [26], it is tempting to speculate that TRIM20 recognises assembled inflammasome complexes to target them for autophagy in order to simultaneously remove the (perceived) threat and the danger signal. In the model, TRIM20 directed autophagy is implemented as dependent on the NLRP3-ASC interaction. Of note, the degradation of parasitophoric vacuoles have been shown to depend on ubiquitylation and interferon-γ (IFN-γ) [57], and IFN-γ has been shown to antagonise NLRP3 inflammasome assembly and signalling [169], supporting a role of K63-linked polyubiquitylation as a switch between NLRP3 dependent signalling and autophagy. It should also be noted that NLRP3 activates autophagy independently of ASC and caspase-1 [4], and the autophagosome targets ubiquitylated NLRP3 [77] – likely via SQSTM1 (p62) [4]. Autophagy constitutes a primitive example of innate immunity [170] (also called xenophagy [171]), and it is known to help clear intracellular pathogens [4]. Taken together, these findings suggests that NLRP3 is targeted for autophagy both before inflammasome assembly – through K63-linked polyubiquitylation – and after inflammasome assembly – through TRIM20-directed precision autophagy.

**Figure 12:**
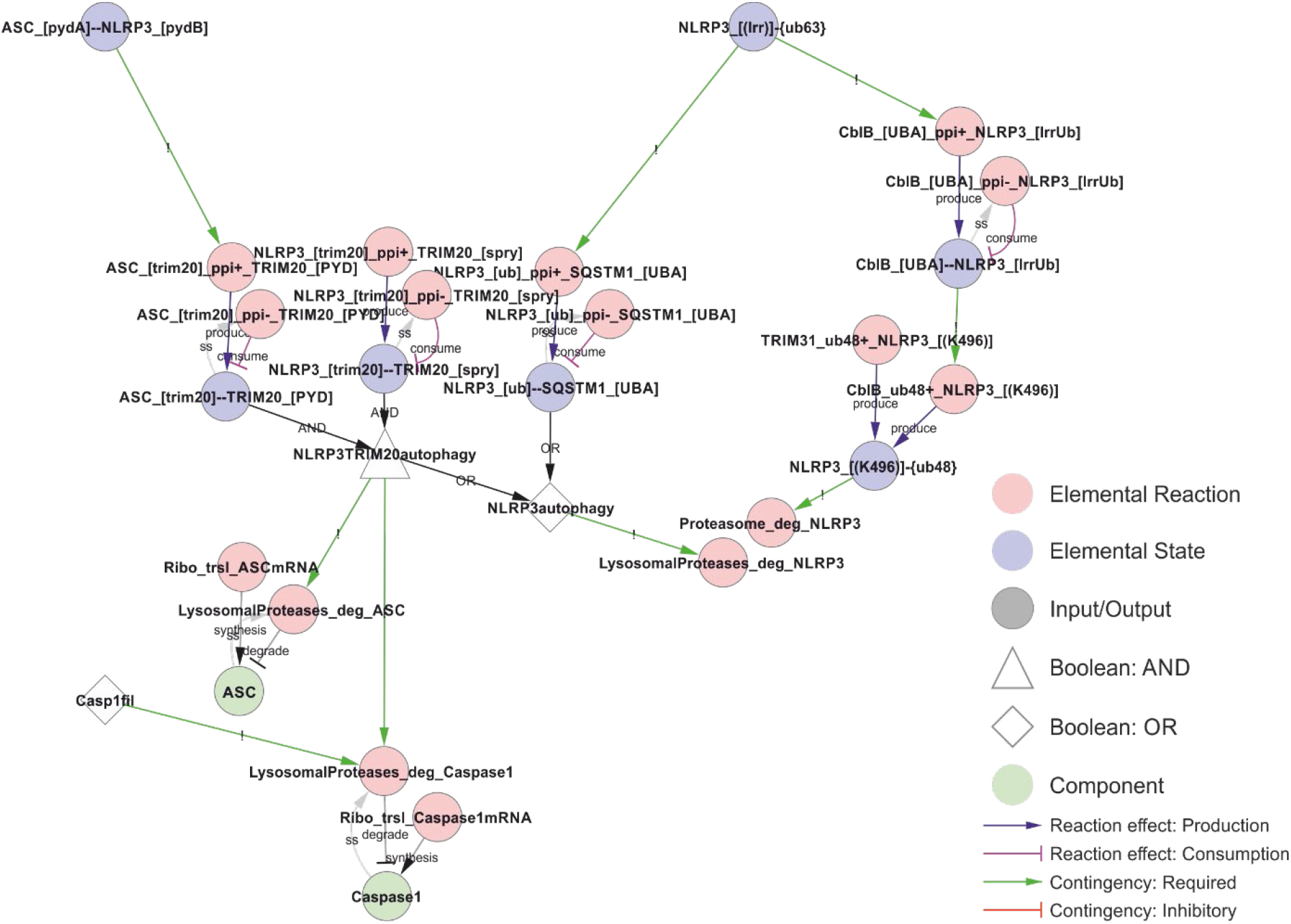
Model of NLRP3 turnover. NLRP3 is degraded both before (right) and after (left) inflammasome assembly. Proteasomal degradation depend on K48-linked ubiquitylation of Lys496, while K63-linked ubiquitin chains are recognised by the autophagy receptor SQSTM1 (p62). In the model, precision autophagy requires inflammasome assembly (interaction between NLRP3 and ASC), which allows simultaneous binding of TRIM20 (pyrin) to both NLRP3 and ASC, leading to autophagosomal degradation of both as well as of caspase-1 if it is part of the complex (Casp1fil). *Walkthrough: Right: K63-linked ubiquitylation of NLRP3’s LRR domain (“NLRP3_[(LRR)]-{ub63}”) allows binding to CblB and SQSTM1. CblB, upon binding (“ClbB_[UBA]--NLRP3_[lrrUb]”), adds K48-linked ubiquitin chains to Lys496 in NLRP3 (“CblB_ub48+_NLRP3_[(K496)]”), targeting NLRP3 for proteasomal degradation (“Proteasome_deg_NLRP3"). The Lys496 site is also targeted constitutively by TRIM31. SQSTM1 binding (“NLRP3_[ub]--SQSTM1_[UBA]”) targets NLRP3 for autophagy and lysosomal degradation (“LysosomalProteases_deg_NLRP3”). In parallel, the ASC-NLRP3 filaments (“ASC_[pydA]-- NLRP3_[pydB]”) are recognised by TRIM20 which binds ASC and NLRP3 simultaneously to target them for autophagy and lysosomal degradation. In case Caspase-1 occurs in the filaments (“Casp1fil”), this is also included in the TRIM20 induced autophagy and degradation*.

### A computational model explaining NLRP3 activation

After completing the network reconstruction, we asked how well the network can explain the known behaviour of the NLRP3 inflammasome system. To answer this question, we generated the bipartite Boolean modelling corresponding to the network and analysed it through simulation (see methods). First, the model was simulated to its natural initial (off) state. Thereafter, we simulated the model from this natural initial state in the presence of LPS, nigericin, or LPS plus nigericin. As can be seen from Figure 13, NLRP3 fails to activate in response to LPS or nigericin *per se*, but does activate in response to LPS plus nigericin, as expected [55]. Similar results were obtained with Pam3csk4 and imiquimod, i.e., neither substance alone triggered NLRP3, but the combination of Pam3csk4 and imiquimod triggered NLRP3 activation, with the difference that K^+^ efflux was a consequence of gasdermin D insertion rather than a trigger in this simulation, and hence occurred only at the end of the simulation. We also mimicked long term / strong LPS exposure by evaluating the effect of cytoplasmic LPS. Here, neither extracellular LPS alone, nor intracellular LPS alone was sufficient to activate NLRP3. However, the combination of intracellular LPS – which triggers non-canonical gasdermin D processing and hence ionic fluxes – and extracellular LPS – which provides a priming and licensing signal – activates NLRP3. Finally, the mitochondrial membrane permeabilisation (MMPT)-triggered cytoplasmic exposure of CL and NLRP3 activation occurred in the presence, but not the absence, of LPS priming. Taken together, the model reproduces basic NLRP3 activation.

**Figure 13:**
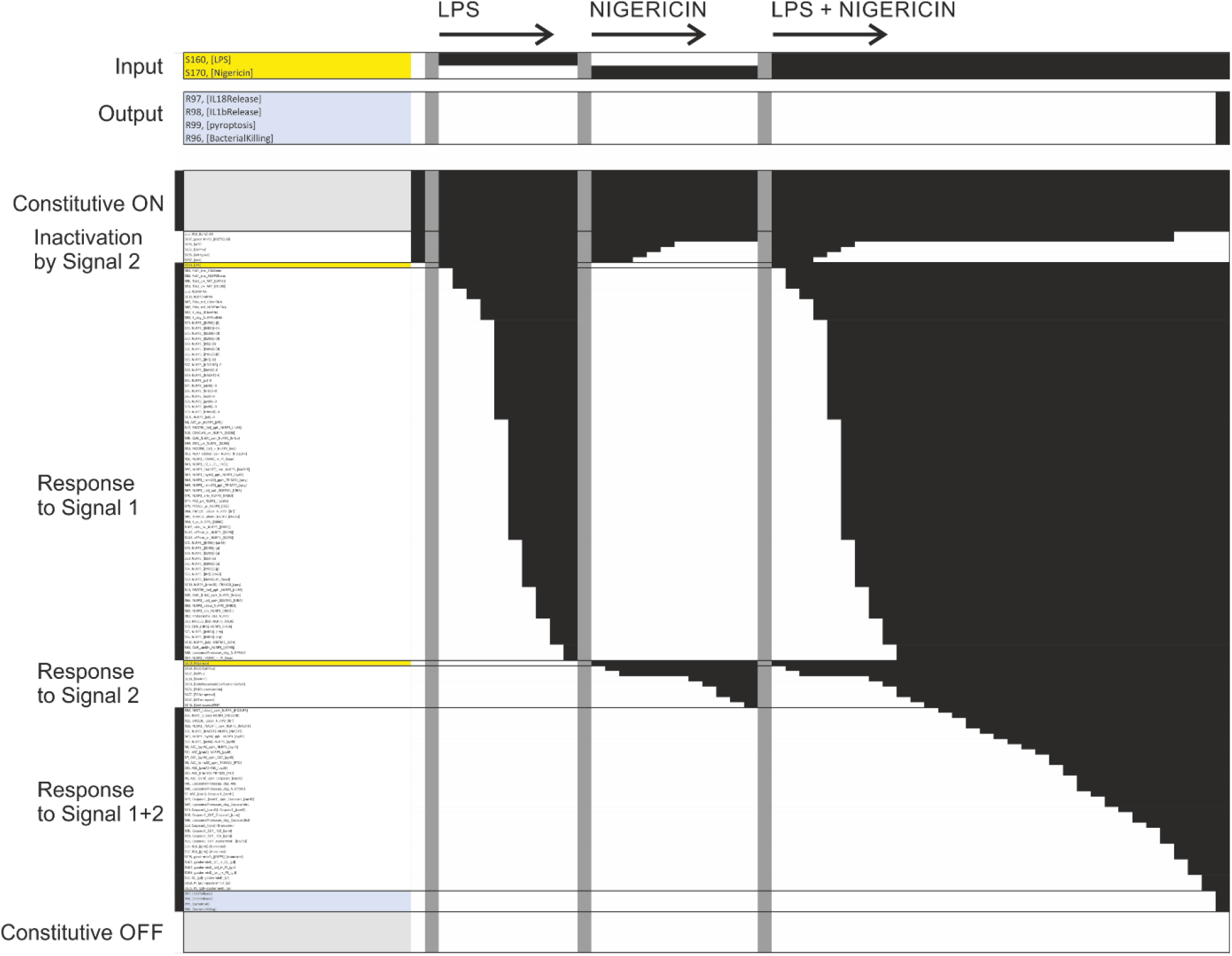
Simulation of the network model. Each row corresponds to one elemental reaction, elemental state, input, output, or component. Each column corresponds to one time point in one of four simulation trajectories. The first trajectory was used to find the natural off state (single column, trajectory not shown). Black squares denote that a particular state was TRUE (ON) at this particular point in the trajectory. Similarly, white squares denote that a particular state was FALSE (OFF). The model was exposed to different stimuli by turning the corresponding input on (TRUE) in the natural off state. Three trajectories are displayed: Exposure to LPS, exposure to nigericin, and exposure to LPS plus nigericin. The input lines are highlighted yellow in the plot and shown enlarged at the very top. Note that exposure to LPS or nigericin only leads to activation of part of the pathway, without triggering the output states (highlighted blue and shown enlarged at the top), while simultaneous exposure to LPS and nigericin leads to inflammasome assembly, IL-1β and IL-18 release, pyroptosis, and, if applicable, bacterial killing. Six blocks of model variables can be recognised in the plot: The uppermost block correspond to constitutively active reactions and states, block two is initially active but inhibited by nigericin, block three includes LPS and all model variables responsive to LPS alone, block four includes nigericin and the model variables activated by nigericin alone, block five includes the model variables responsive to LPS and nigericin but neither alone, and the final block includes the variables that remain off under all simulation conditions shown here. The last include other inputs (including mutations and chemical perturbations) not used in the simulation, and reactions and states directly downstream of those. The different trajectories are separated by vertical grey bars. The model variables (rows) have been sorted on the sequence of activation, making the sequential activation especially after LPS plus nigericin treatment clear.

Using the model, we evaluated the effect of inhibitors on NLRP3 activation. First, we tested MCC950, which initially failed to prevent NLRP3 activation in response to nigericin and LPS. It turned out that the binding of ATP to newly synthesised NLRP3 sufficed to bypass MCC950 inhibition. When MCC950 inhibit both ADP-to-ATP exchange and ATP binding to empty NLRP3, the activation is interrupted after NEK7 binding but before any downstream events, imposing a complete inhibition of NLRP3 activation. Second, we analysed the effect of BRCC36 inhibition by the general deubiquitinase inhibitor G5 [61]. In the model, G5 completely inhibits deubiquitylation of NLRP3 LRR domain, but fail to prevent NLRP3 activation due to priming dependent synthesis of new (unubiquitylated) NLRP3. Third, we analysed the effect of the PP2Aca inhibitor Okadaic Acid (OKA) [58]. In the model, OKA completely inhibits the dephosphorylation of NLRP3 at Ser5, but again the lack of dephosphorylation is offset by synthesis of new (unphosphorylated) NLRP3. If the bypass of G5 and OKA inhibition by protein synthesis is relevant in vivo remains unclear, but it demonstrates the limitation of negative licensing, and may help illustrate why overexpression of NLRP3 can make NLRP3 activation independent of Signal 1.

Furthermore, we implemented and tested two CAPS mutations and a truncated version of NLRP3 lacking the LRR domain in the model. First, NLRP3 D305G is implemented as an inhibitor of PKA-mediated phosphorylation of Ser295. This mutation failed to activate NLRP3 alone or in combination with LPS in the simulations, suggesting that the clinical symptoms may be due to quantitative modulation of the ATP cycle that this qualitative model cannot capture. Second, in the model, the NLRP3 mutation T346M stabilises the open conformation, making activation of NLRP3 ATP-binding independent, effectively bypassing the need for PI(4)P binding to achieve the open structure in NLRP3 in the model. This alone is not enough to activate NLRP3, not even in the presence of LPS. However, this is due to the model requirement of PI(4)P localisation to the centrosome, which should not be needed if NLRP3 activation is PI(4)P binding independent. If the model accounts for this, then NLRP3 T346M is indeed activated upon priming with LPS alone. Finally, we tested the effect of deleting the complete LRR domain, mimicking the “miniNLRP3” experiments [38]. This is implemented by inhibiting all reactions involving exclusively the LRR domain (table 2). The truncated NLRP3 phenocopies the full length NLRP3 in the model for PI(4)P dependent activation, i.e., it does not respond to LPS or nigericin alone, but it is activated by LPS and nigericin together. However, miniNLRP3 failed to respond to MMPT and cytoplasmic CL exposure in the model, as NLRP3 binds CL via its LRR domain, suggesting that miniNLRP3 should not be activatable by the CL-axis *in vivo*.

**Table 2:**
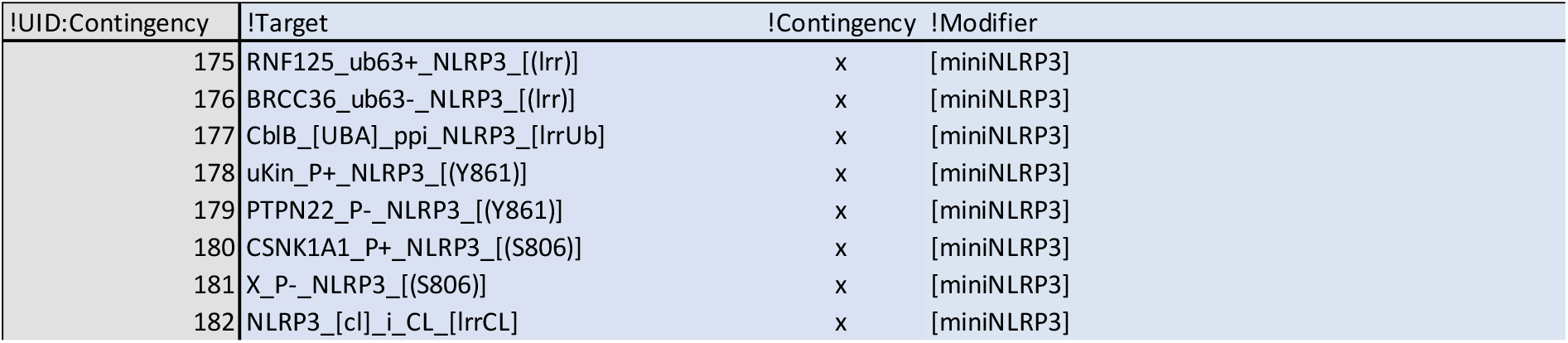
Implementation of miniNLRP3. MiniNLRP3 is a truncated form of NLRP3 completely lacking the LRR domain, and hence all reactions targeting this domain is unavailable. This is implemented as an input “[miniNLRP3]” that inhibits all reactions targeting this domain.

## Discussion

The trigger signal for NLRP3 remains an open question. In this work, we have made the Ansatz that all NLRP3 trigger signals converge on one common cellular perturbation, and that this perturbation trigger NLRP3 activation – given that the priming and licensing conditions are fulfilled. We find that the evidence supports this, at least when it comes to the activation along the PI(4)P-NEK7 axis as all trigger signals lead to accumulation of mobile intracellular vesicles, that accumulate PI(4)P, and which therefore, at least in principle, can support NLRP3 activation. There are some key studies that support this notion. Most importantly, the PI(4)P binding region is sufficient to impose NLRP3-like regulation to all three types of stimuli to NLRP6, which normally does not respond to those stimuli [18]. This localises the Signal 2-sensing to this region, and PI(4)P binding is the only regulatory feature that has been mapped to this region, strongly suggesting that this is the critical regulatory input. Second, PI(4)P binding is in itself not enough [31], and NLRP3 release from its resting position on PI(4)P-containing Golgi membranes is necessary for activation [16], as is microtubule-based transport [127]. However, some membrane dispersing toxins (shown for monensin) do not result in NLRP3 activation [140], showing that also dispersal is insufficient for activation, suggesting that either PI(4)P accumulation or microtubule transport is regulated. At least the first is supported by previous data, as LMP has been found to trigger rapid recruitment of PI4K after lysosomal rupture/depolarisation [163]. This suggest that the common feature of NLRP3 regulation is most likely osmotic lysis and/or depolarisation of internal vesicles, which recruits PI4K and can be transported to the centrosome to create the conditions for NEK7 and PI(4)P-dependent activation of NLRP3.

Taking one step back, to the question how the diverse NLRP3 triggers cause osmotic lysis or depolarisation, we propose that the ability of the cell to maintain the ion gradient against leakage is critically disrupted by energy depletion (which directly affects the ion pumps) and/or membrane permeabilization. This would immediately result in a loss of osmotic integrity [152], and direct inhibition of both the plasma membrane (Na^+^/K^+^-ATPase; [157]) and vacuolar (V-ATPase; [149]) pumps have indeed been found to trigger NLRP3 activation after priming. The findings that hypo-osmolarity can trigger NLRP3 and hyper-osmolarity can suppress NLRP3 activation by other triggers [123] support this notion. However, there are also evidence for a completely different axis of NLRP3 activation by which NLRP3 can be activated by CL binding to the c-terminal LRR domain [66], which lies in the opposite end of the protein to the n-terminal PI(4)P binding domain. Moreover, CL has been shown to recruit caspase-8 [128], which is an essential component of the NEK7-independent NLRP3 activation [78]. CL in the cytoplasm could indicate a bacterial infection that should be directly targeted for destruction, suggesting that the normal time delay in inflammasome activation may be undesirable. Hence, it is possible that there is a CL-caspase-8 axis of NLRP3 activation that is fundamentally different – from structure of complex formation to regulation by post-translational modifications – than the PI(4)P-NEK7 axis of NLRP3 activation. By design or default, almost all the data we have encountered seem to have studied the latter, as a “mini-NLRP3” lacking the LRR domain than bind CL reproduces virtually all known regulation [38]. Hence, the potential CL-caspase-8 axis appears largely unexplored and its mechanistic architecture and relevance for NLRP3 activation in health and disease is therefore currently difficult to establish.

The network reconstruction process has highlighted the multiple roles of NLRP3: As a signalling platform, as an activator of autophagy, and as a mediator of direct bactericidal action. The overlap in lipid affinity between NLRP3 and gasdermin D is striking [25, 31, 66], and this, together with the report that gasdermin D is part of the NLRP3 inflammasome complex [67], suggests that the NLRP3 inflammasome may catalyse the targeted insertion of bactericidal gasdermin D pores into the membrane on which it is activated [5, 30]. At the same time, NLRP3 can activate autophagy/xenophagy [170, 171] to help clear pathogens [4]. However, pathogens are also known to subvert intracellular organelles to form replicative niches [172], e.g., by preventing phagosomal-lysosomal fusion [173], which suggests a potential overlap between such subverted compartments and the compartments where NLRP3 activation may occur. Hence, NLRP3 may be able to recognise intracellular pathogens both directly (through CL binding) and through perturbation of the membrane of intracellular compartments, coordinating a membrane attack or at least permeabilisation of the infected compartment with autophagic disposal and intercellular signalling through IL-1β and IL-18. With limited insult and successful clearance, a targeted insertion of gasdermin D should leave the cell able to recover and there are observations in the literature showing that IL-1β release and Gasdermin D pores are possible without the presence of cell death markers [174, 175]. However, if insults are saturated, an extensive gasdermin D insertion may trigger the positive feedback loop through ion leakage, leading to irreversible NLRP3 activation. In fact, pyroptosis may well be an emergency response to unmanageable infection or damage, or an accidental side effect of a protective and essentially homeostatic process. Such a harmful outcome may explain the extensive licensing regulation of NLRP3, which appears to be only partially dependent on Signal 1, and which may serve to restrict NLRP3 activation to valid target compartments and to selectively exclude NLRP3 activation and hence gasdermin D insertion from, e.g., the plasma membrane, where at least extensive insertion would likely be suicidal. However, if gasdermin D insertion in the plasma membrane is prevented, it leaves the question as to how IL-1β and IL-18 are released during physiological responses. Of note, it was found that NLRP3 activation triggers shedding of IL-1β and IL-18 containing exosomes [176], which are exported to the extracellular space where they may release their cytokines through gasdermin D pores or vesicular lysis without impacting cell integrity. It is tempting to speculate that the physiological NLRP3 response leads to targeted insertion of gasdermin D into specific vesicles, that are selectively loaded with locally processed IL-1β and IL-18, engulfed through autophagy and delivered to the extracellular space through exocytosis. In any case, NLRP3 has been shown to be a critical regulator of intracellular defence and intercellular signalling.

The model presented here is essentially a model of PI(4)P-NEK7 dependent activation of NLRP3, and even this is merely a snapshot based on the currently available data and knowledge. Moreover, it was not possible to cover even the already available literature in the field, as a search for “NLRP3” alone on PubMed yields more than 15,000 hits. This highlights the need to build a formal reusable knowledge base that the community can use, update, and expand as the field progresses. It is important that such a knowledge base is highly composable - i.e., allows statements to be added, edited, or removed individually, and arbitrary parts to be extracted and/or combined for analysis – to allow the distributed work necessary for sustainable community efforts and to make it useful for a wide range of projects. To this end, the mechanistic knowledge of the NLRP3 system is broken down into minimal statements – elemental reactions and contingencies – which are defined in terms of site-specific elemental states. The advantage of this approach is that the knowledge of individual reactions can be formulated independently, including both the effect (the elemental reaction) and the regulation (the contingencies), so that these statements can be individually evaluated, modified, and added or removed. However, it also requires this information to be available in the literature, i.e., that the effect of specific modifications on a specific reaction has been examined directly, which is not always the case. Here, we use targeted literature searches to establish such a mechanistic network for the core NLRP3 regulation including some, but not all, reported modification sites and interaction partners, as we have been unable to find sufficient mechanistic data for several of the components and modifications suggested in the literature to be of importance for NLRP3 regulation. We focussed on generating a consistent model that could explain NLRP3 activation from the existing data, rather than on highlighting inconsistencies, meaning that there are a number of assumptions present in the model. At this level, the network is effectively a formal and highly reusable literature review, with the added feature that all statements must be precise and internally consistent, and this curation and formalisation process is indeed the most challenging part of building a network model. However, once this knowledge base (consisting of elemental reactions and contingencies) is compiled, it enables visualisation and computational analysis of (selected parts of) the complete knowledge base. Here, we make use of the rxncon regulatory graph to visualise the causal information flow through the network, to make the regulatory structure of the network accessible to readers. Moreover, the biological knowledge base can be automatically converted into a bipartite Boolean model (bBM). The limitation of the bBM is that it can only make qualitative predictions (yes/no, active/inactive), without quantities and meaningful time resolution. With that said, it is uniquely defined by the biological knowledge base, does not need parametrisation or model optimisation, and can hence be used to directly evaluate the knowledge base. Here, we use it to evaluate if the assembled knowledge suffices to explain the known system regulation (does it respond to the given input(s)?), and if it can predict the effect of inhibitors and mutations (how is the response altered by a given combination of inhibitors and/or mutations?). The network does indeed suffice to explain NLRP3 activation to a range of inputs, although the effect of inhibitors and mutations are sometimes less clear to evaluate with the bBM. The simulation results suggests that, given significant *de novo* synthesis of NLRP3, the negative licensing may be ineffectual. However, it is also clear that this modelling scheme cannot explain quantitative effects, and the importance of quantitative effects may be a general feature in NLRP3 activation, including at the level of synthesis, spatiotemporal restriction, and regulation by autophagic degradation. Despite these limitations, the knowledge base of molecular mechanisms presented here is an internally consistent knowledge base that contain the mechanisms that are necessary and sufficient to explain the qualitative behaviour of the core NLRP3 network, which can be processed and analysed computationally, and which can easily be adapted and extended by the community as new data and knowledge become available.

## Supporting information

Figure S1 - high resolution of Figure 1

Supplementary Table 1 - rxncon model

## Acknowledgements

We would like to thank Dr. Alexander Persson for many useful discussions on the NLRP3 inflammasome and to acknowledge scientific support from the Exploring Inflammation in Health and Disease (X-HiDE) Consortium, which is a strategic research profile at Örebro University funded by the Knowledge Foundation (20200017).

