## Supplementary figures and images for "A detailed Molecular Network Map and Model of the NLRP3 Inflammasome"

### Figure S1 - high resolution of Figure 1

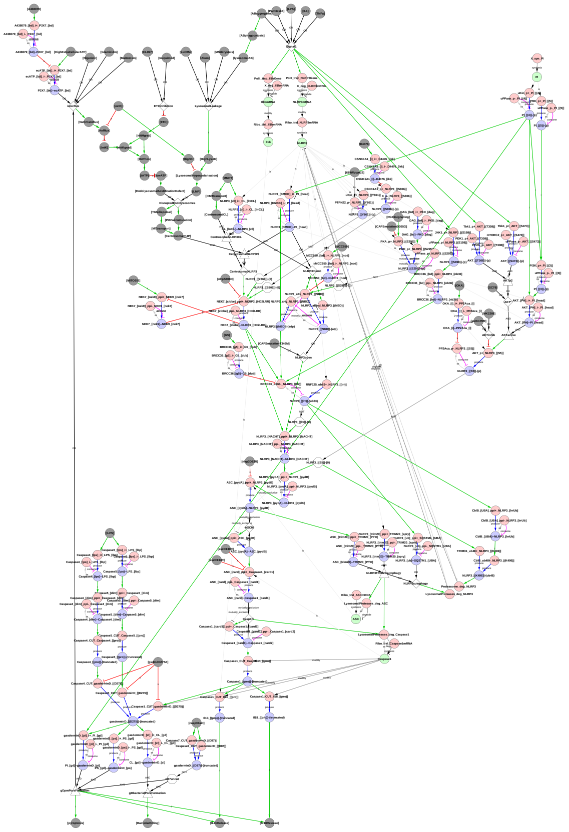
